# A novel effect of the antigenic peptides position for presentation upon MHC class I

**DOI:** 10.1101/653170

**Authors:** Jun Imai, Mayu Otani, Eri Koike, Yuuki Yokoyama, Mikako Maruya, Shigeo Koyasu, Takahiro Sakai

**Affiliations:** Laboratory of Physiological Chemistry, Faculty of Pharmacy, Takasaki University of Health and Welfare, 60 Nakaorui-machi, Takasaki-shi, Gunma 370-0033, Japan; Keio Research Park, Keio University School of Medicine, 35 Shinano-machi, Shinjuku-ku, Tokyo 160-8582, Japan; Department of Microbiology and Immunology, Keio University School of Medicine, 35 Shinano-machi, Shinjuku-ku, Tokyo 160-8582, Japan; CREST, Japan Science and Technology Corporation, Kawaguchi-shi, Saitama 332-0012, Japan; Center for Integrative Medical Sciences, RIKEN Yokohama Institute, Yokohama, Kanagawa 230-0045, Japan; Laboratory for Immune Cell Systems, RIKEN Centre for Integrative Medical Sciences (IMS), Yokohama, Kanagawa 230-0045, JAPAN

**Keywords:** antigenic peptide, *de novo* proteins synthesis, direct presentation, DRiPs, IFN-γ

## Abstract

MHC-I molecules are expressed on the cell surface complexed with oligopeptides, most of which are generated from intracellular proteins by the ubiquitin-proteasome pathways and would be roughly proportional to the relative abundance of proteins and their rate of degradation. Thus MHC-I together with peptides function as immunological self-markers to exhibit the information about the repertoire of proteins expressed in a given cell to the immune system, and this process is called antigen (Ag) direct-presentation. Herein, we report a novel rule for the preference of peptides selected for the direct presentation; the N-terminally located antigenic peptides are more efficiently complexed with MHC-I than the C-terminally located peptides on the same protein. The superiority is largely dependent upon *de novo* proteins synthesis, degradation by proteasomes, and less dependent upon stabilities of proteins, indicating that this difference derived from rapidly degraded newly synthesized proteins such as defective ribosomal products (DRiPs). The effects of those N-terminal predominance was comparable with the enhanced MHC-I presentation by IFN-γ suggesting that they might play important roles in the adaptive immunity.

## Introduction

MHC-I molecules are expressed on the cell surface complexed with antigenic peptide, typically 8-11 residues long, most of which were generated from cytosolic proteins by the ubiquitin-proteasome pathways (Cresswell et al., 2005; Groothuis and Neefjes, 2005; Loureiro and Ploegh 2006; Shastri et al., 2005; Trombetta and Mellman 2005; Yewdell et al., 2003). The repertoire of antigenic peptide would be roughly proportional to the relative abundance of proteins and their rate of degradation. Antigenic peptides together with MHC-I function as immunological self-markers to exhibit the information about the repertoire of proteins expressed in a given cell to the immune system and this process is called antigen (Ag) direct-presentation (Janeway et al., 2001). Cancer or virally infected cells express disease-specific non-self proteins thus display non-self peptides upon MHC-I (Janeway et al., 2001). Cytotoxic T lymphocytes (CTL) can detect the non-self peptides with MHC-I and lyse target cells (Janeway et al., 2001).

There have been several numbers of databases for MHC-I-binding peptides (The Immune Epitope Database; hrrp://www.iedb.org/home_v3.php, etc.). Also, Ag-peptides are predictable for some MHC alleles, dependent upon the binding motif found in the given Ags for MHC-I (SYFPEITHI; http://www.syfpeithi.de/scripts/MHCServer.dll/home.htm, etc.). It is also likely that these peptides are generated by the cellular protein quality control systems. Misfolded proteins and non-functional polypeptides (Starck et al., 2016) are a possible source of Ag-peptides in addition to once correctly folded proteins. Accumulated evidence shows that considerable amounts of Ag-peptides are derived from polypeptides so-called defective ribosomal products (DRiPs); polypeptides arising from prematurely terminated proteins and misfolded proteins derived from unsuccessful translation (Anton and Yewdell, 2014; Apcher et al., 2011; Apcher et al., 2012; Apcher et al., 2013; David et al., 2012; Dolan et al., 2011; Eisenlohr et al., 2007; Khan et al., 2001; Princiotta et al., 2003; Qian, Princiottam, et al., 2006; Qian Reits et al., 2006; Reits et al., 2000; Rock et al., 2014; Schlosser et al., 2007; Schubert et al., 2000; Starck and Shastri 2011; Voo et al., 2004; Yewdell et al., 2001; Yewdell and Nicchitta, 2006). Because of its fate of fast degradation, DRiPs enables the rapid presentation of Ags to CTL. Given the mechanisms of the cellular protein quality control system, the repertoire of peptides generated from these two sources, mature proteins and DRiPs, would not be the same, because DRiPs contain truncated and premature forms of proteins. The relative share of these two sources for Ag-peptide generation is not entirely determined (Bourdetsky et al., 2014; Rock et al., 2014; Wei et al., 2015). The prospective difference of repertoire of peptides among these sources prompted us to investigate the rule for the preference for Ag-peptides. We have devised methods to detect a quantitative correlation between the expression of pOV8 harboring proteins and the generation of a pOV8/MHC-I complex. We found a new rule for the preference of Ag-peptides; the N-terminally located Ag-peptide is more efficiently complexed with MHC-I than the C-terminally located peptide from the same Ag. The advance of the N-terminal predominance was considerably equivalent with the effect of IFN-γ upon MHC-I presentation suggesting that this rule might play important roles in the adaptive immunity.

## Results

### The presentation efficiency of endogenous pOV8 is dependent upon its location on the protein

We investigated the relationship between the expression of a given protein and the presentation of an antigenic peptide derived from the protein using a panel of chimeric fluorescence proteins (Fig. 1*A, F*, and *G*) with the pOV8 epitope (*SIINFEKL*). The pOV8 epitope was flanked by five flanking amino acids each in its N- and C-terminus (LEQLE*SIINFEKL*TEWTS) (Qian et al., 2002). We examined the level of protein expression and the amount of peptide-MHC-I complex by flow cytometry with the 25-D1.16 monoclonal antibody, which recognizes pOV8 epitope complexed with the H-2kb allele of MHC class I (pOV8/Kb complex) (Porgador et al., 1997). In all chimeric fluorescence proteins except purified His-tagged proteins (Fig. 5) used in this study, the second amino acid was standardized by Val, so as to avoid the difference of stabilities for chimeric fluorescence proteins caused by the N-terminal rule (the second amino acid of His-tagged proteins were normalized as Ala by vector sequence). First, we characterized the amounts of the pOV8/Kb complex as the function of the expression level of chimeric fluorescent proteins using a dendritic cell (DC)-like cell line DC2.4, a thymoma cell line EL4, and a fibrosarcoma cell line MC57G. 25-D1.16 staining was detected specifically in cells that transfected with chimeric fluorescent proteins with pOV8 but not in cells transfected with only fluorescent proteins (Fig. 1*A, G*). On the contrary, fluorescence was detected only in cells transfected with fluorescent proteins but not in cells transfected with OVA alone (Fig. 1*F*), indicating that this method completed our intention. From the overlapping fluorescence histograms for transfectants of the pair of constructs (color combine), we can compare the amounts of pOV8/Kb derived from the construct with pOV8 on the N-terminal (colored in red) with those from the construct with pOV8 on the C-terminal (colored in green) of the same protein. Since the expression level of each chimeric protein was quantified by its fluorescence intensity, it was clear that the efficiency of pOV8/Kb complex formation was higher in the N-terminal pOV8 than the C-terminal pOV8 (Fig. 1*A, F, G* right-most). Next, we examined the correlational functions among the fluorescent intensity and the expression level of the pair of fluorescent chimeric proteins, pOV8-YFP and YFP-pOV8, by flow cytometry with the monoclonal anti-GFP antibody clone E4 and RQ2. From the overlapping fluorescence histograms as Fig. 1A, the correlation was approximately equivalent in DC2.4 (Fig. 1*B*). We also checked these differences were neither the results of the difference in transcription (Fig. 1*C*), nor expression (Fig. 1*D*), nor the stabilities (Fig. 1*E*), nor the localizations (Fig. 1*H*), and nor the fluorescence intensities (Fig. 5*B*) of the pair of fluorescent chimeric proteins. It was noteworthy that 4-6 h was enough to bring about the N-terminal predominance from the subsequent experiments (Fig. 3*B, C*, and *D*), stabilities of pOV8-YFP and YFP-pOV8 were equivalents after 6 h of treatments by Cycloheximide (Fig. 1*E*), by α-amanitin (Fig. 4*F*). It was also notable that the N-terminal predominance was detectable with Azamigreen (Fig. 1*G*).

**Figure 1.**
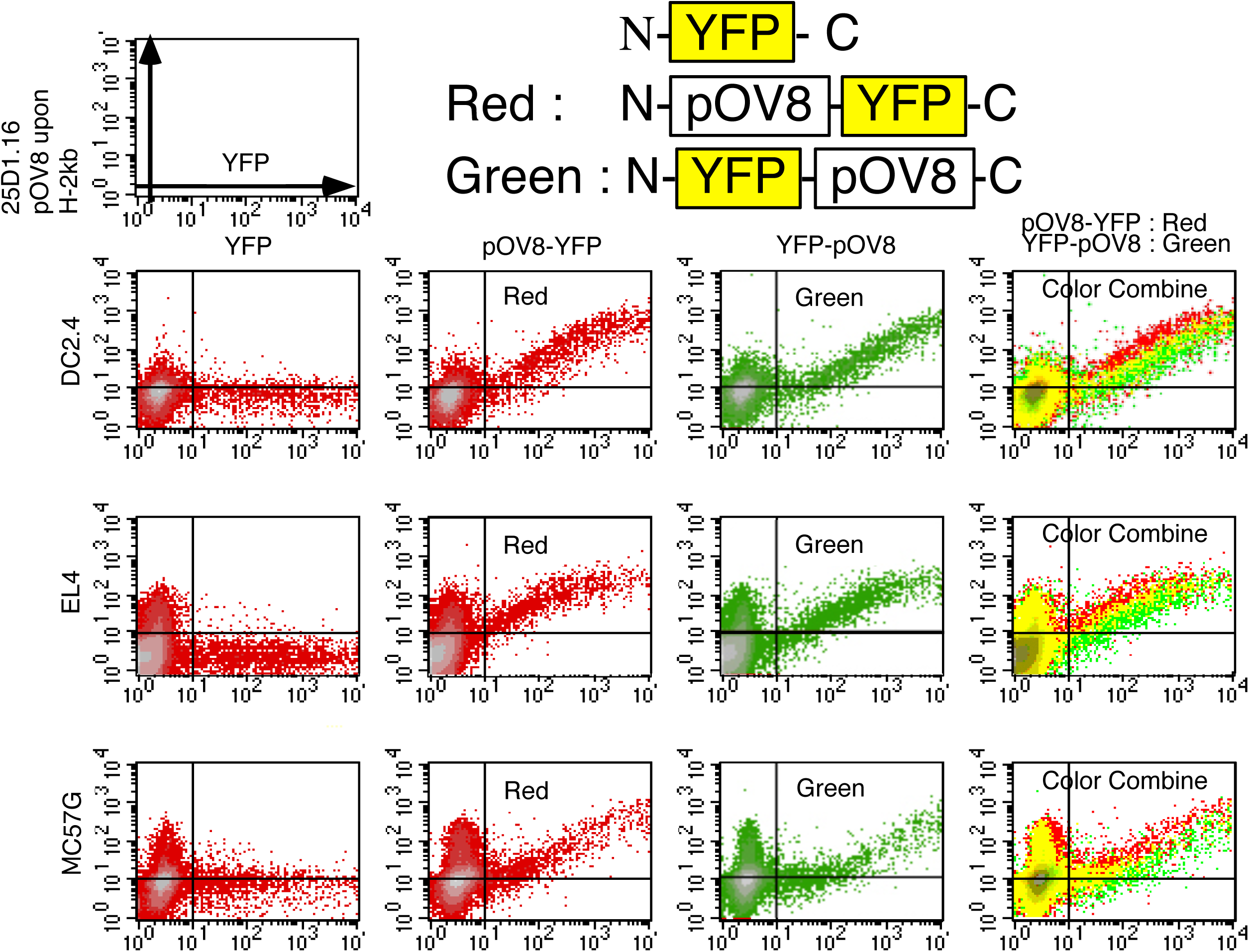
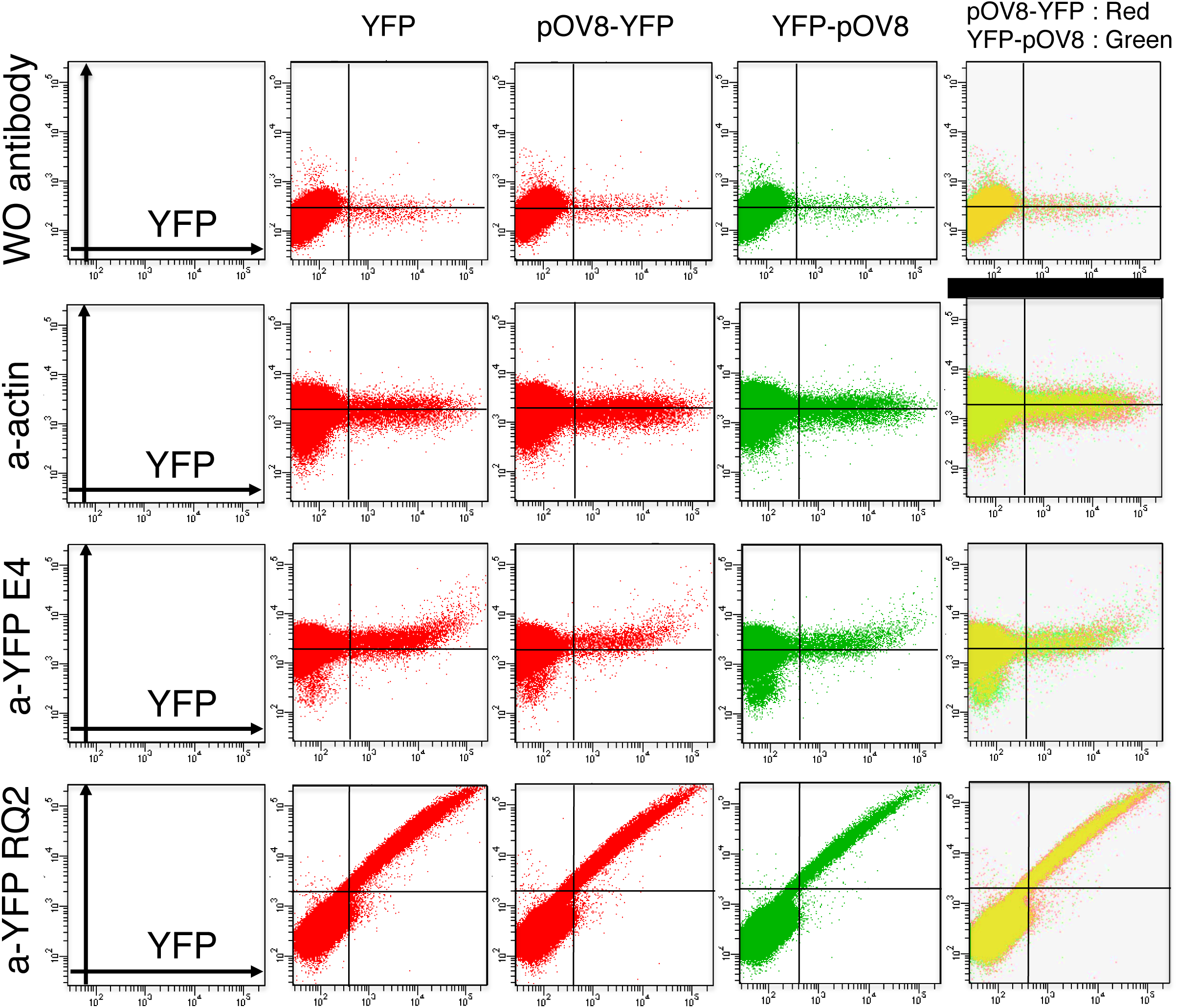
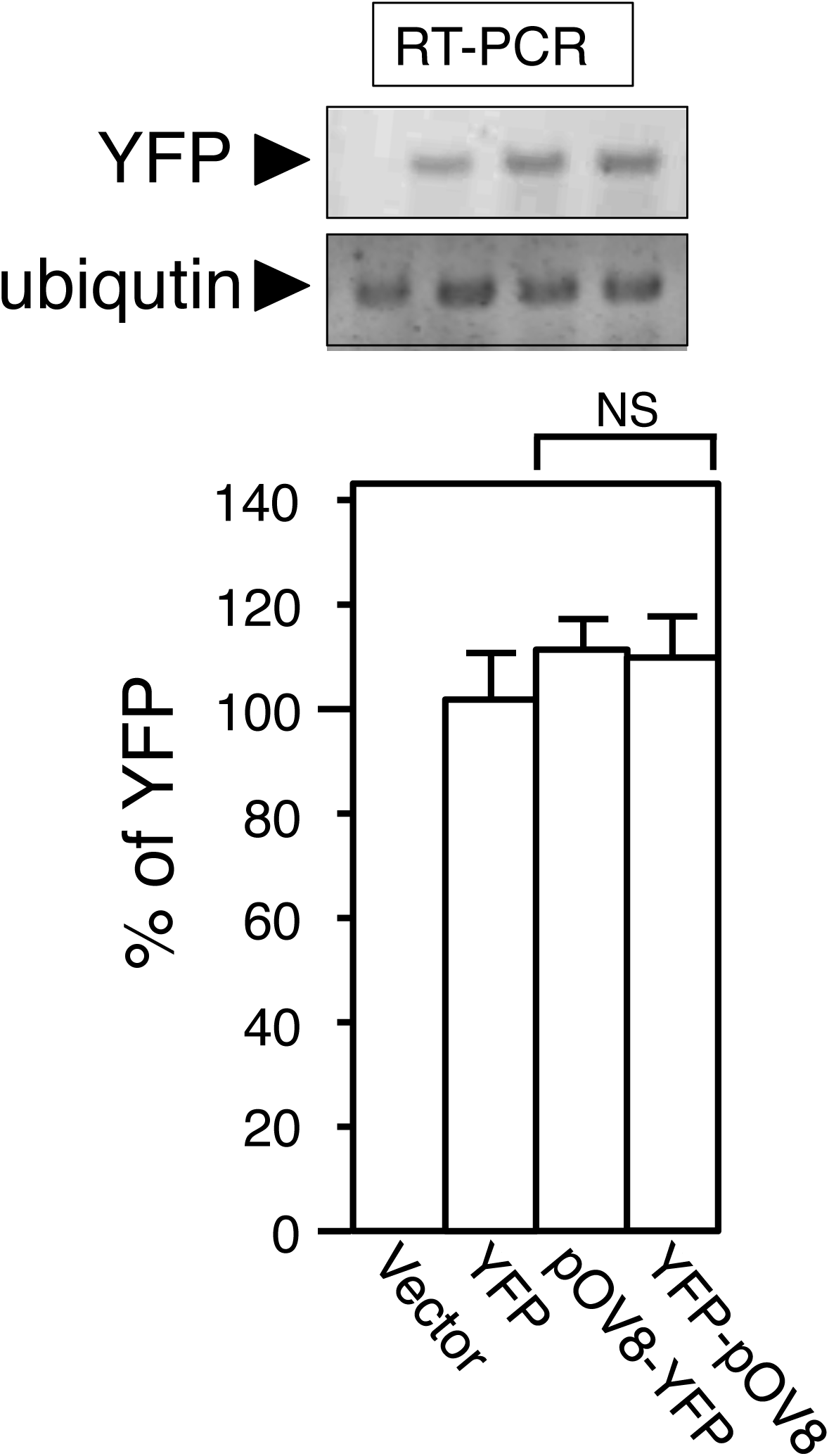
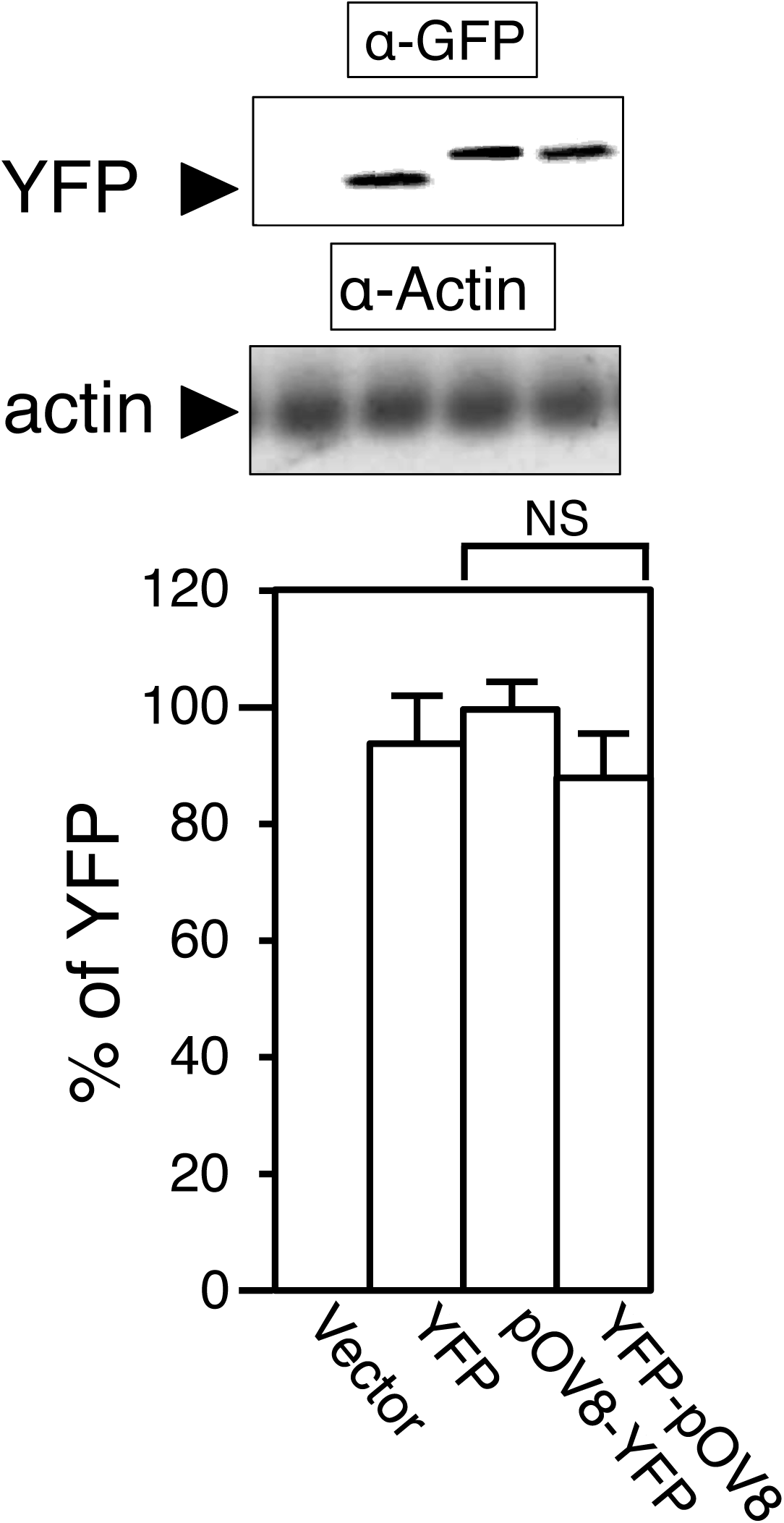
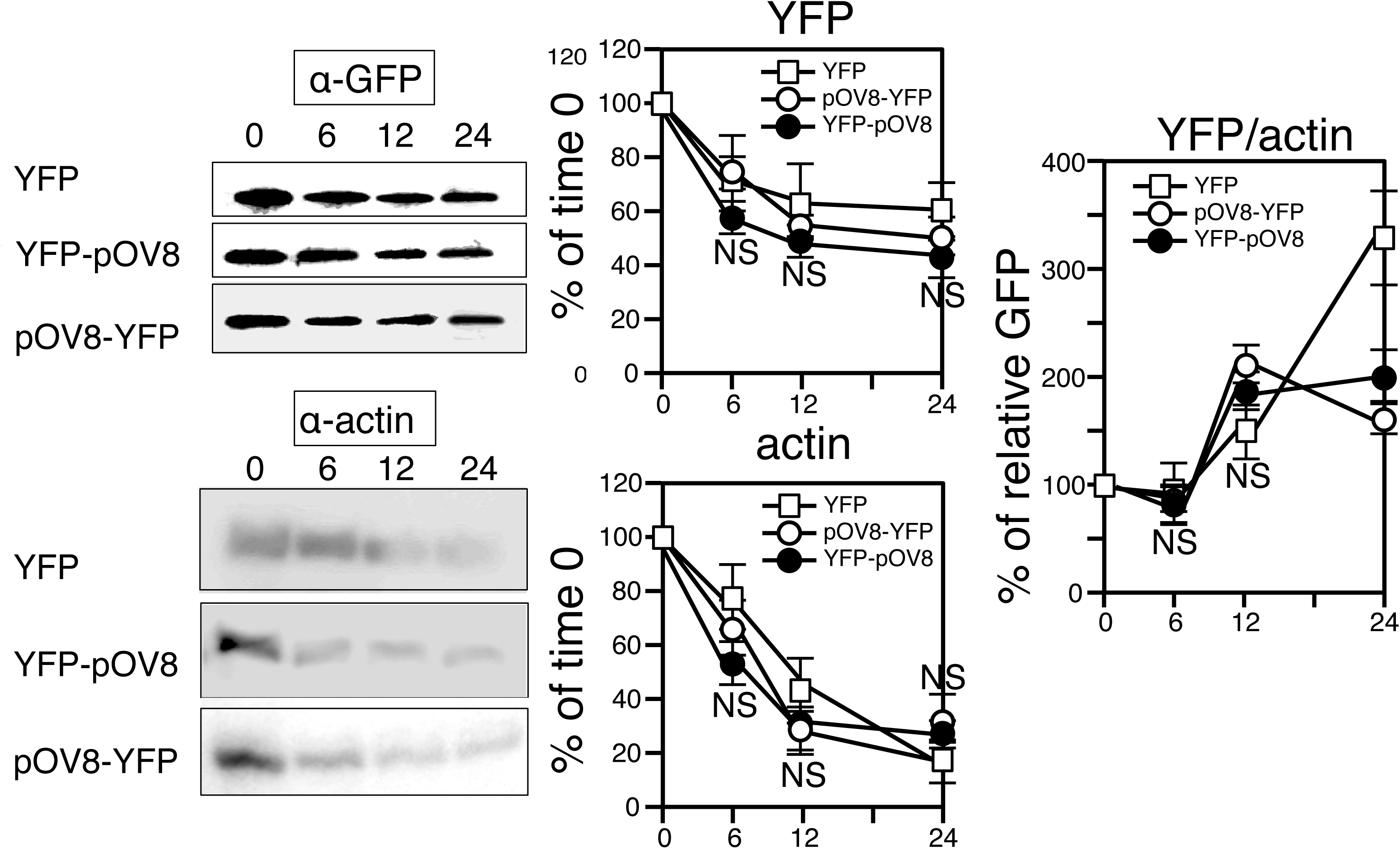
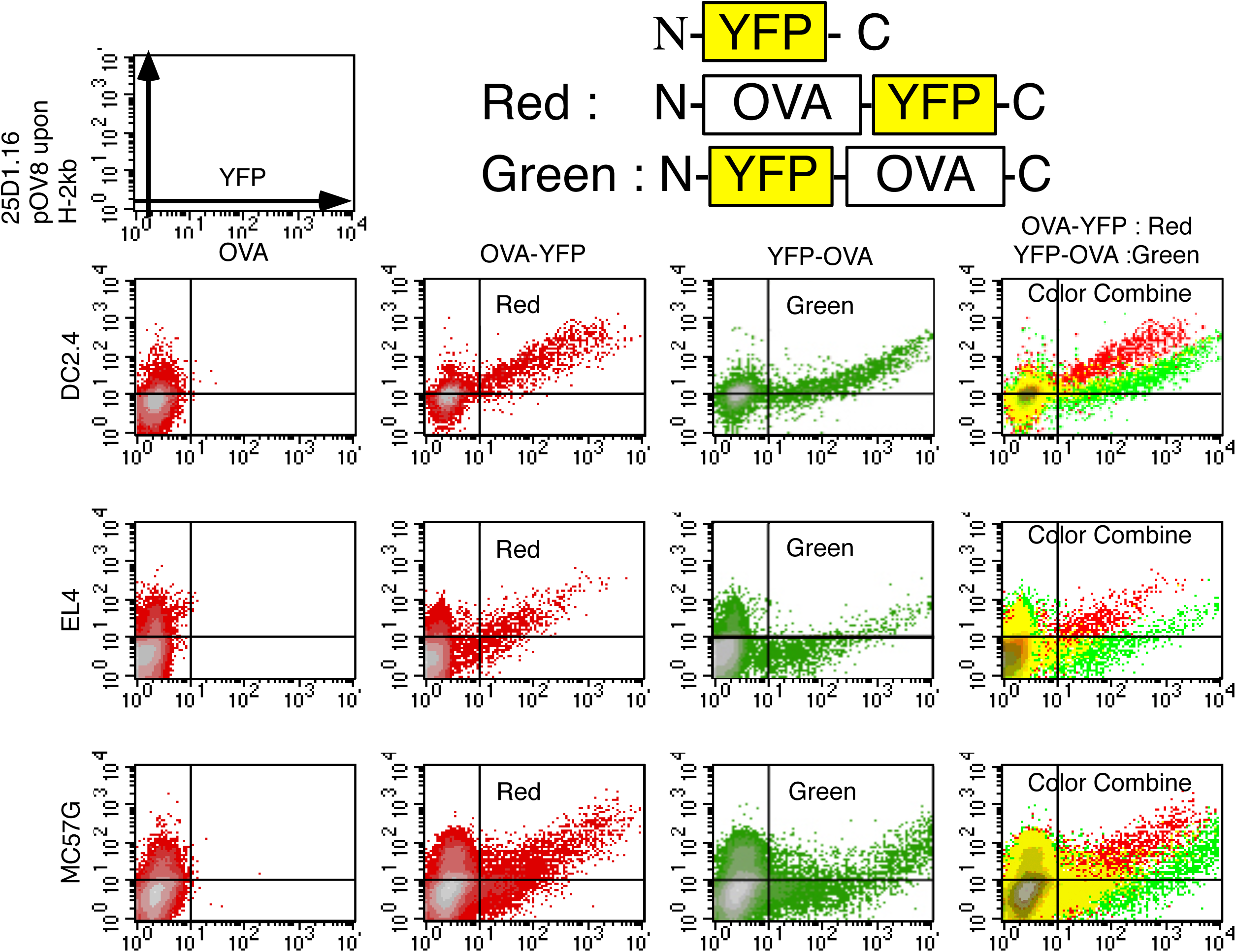
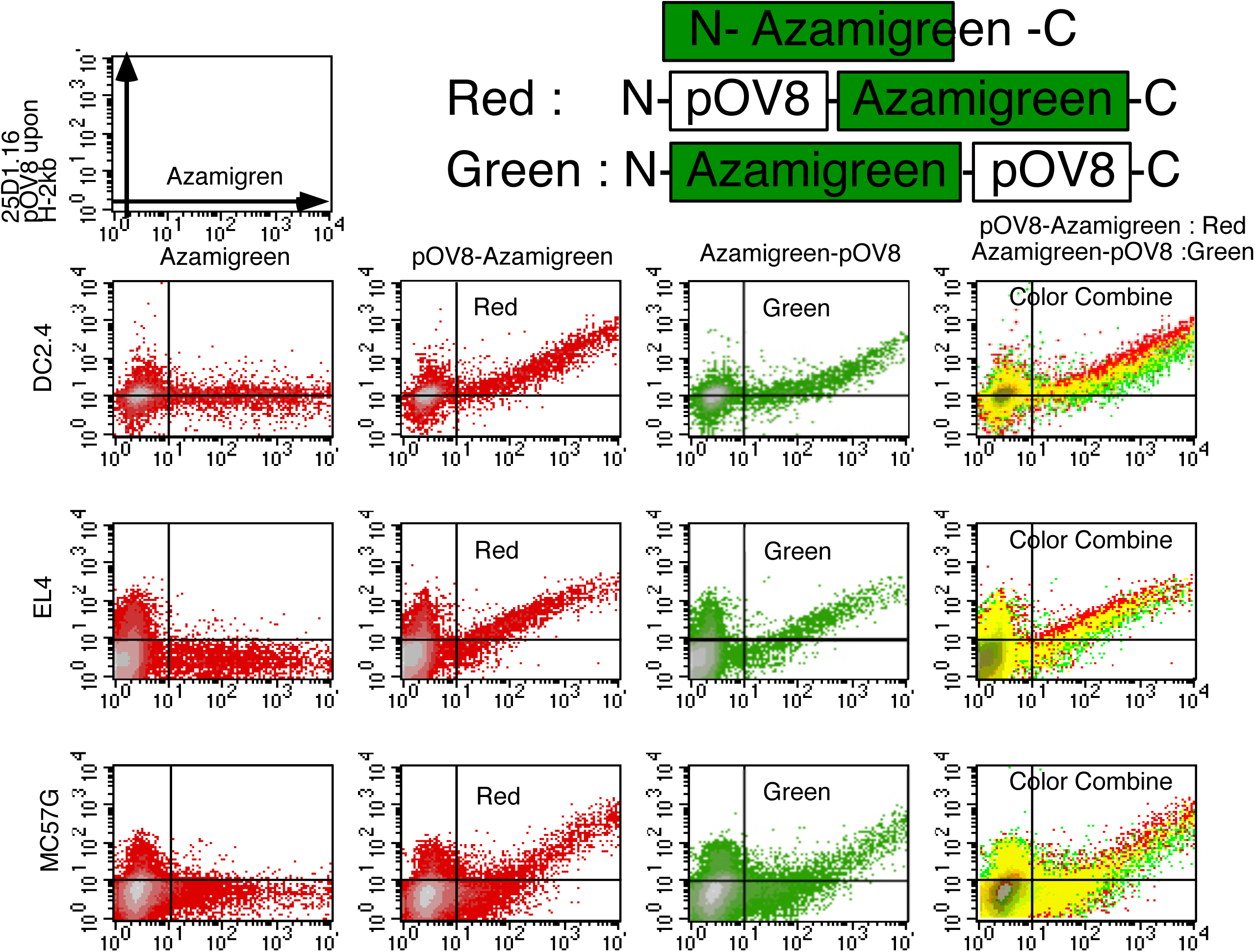
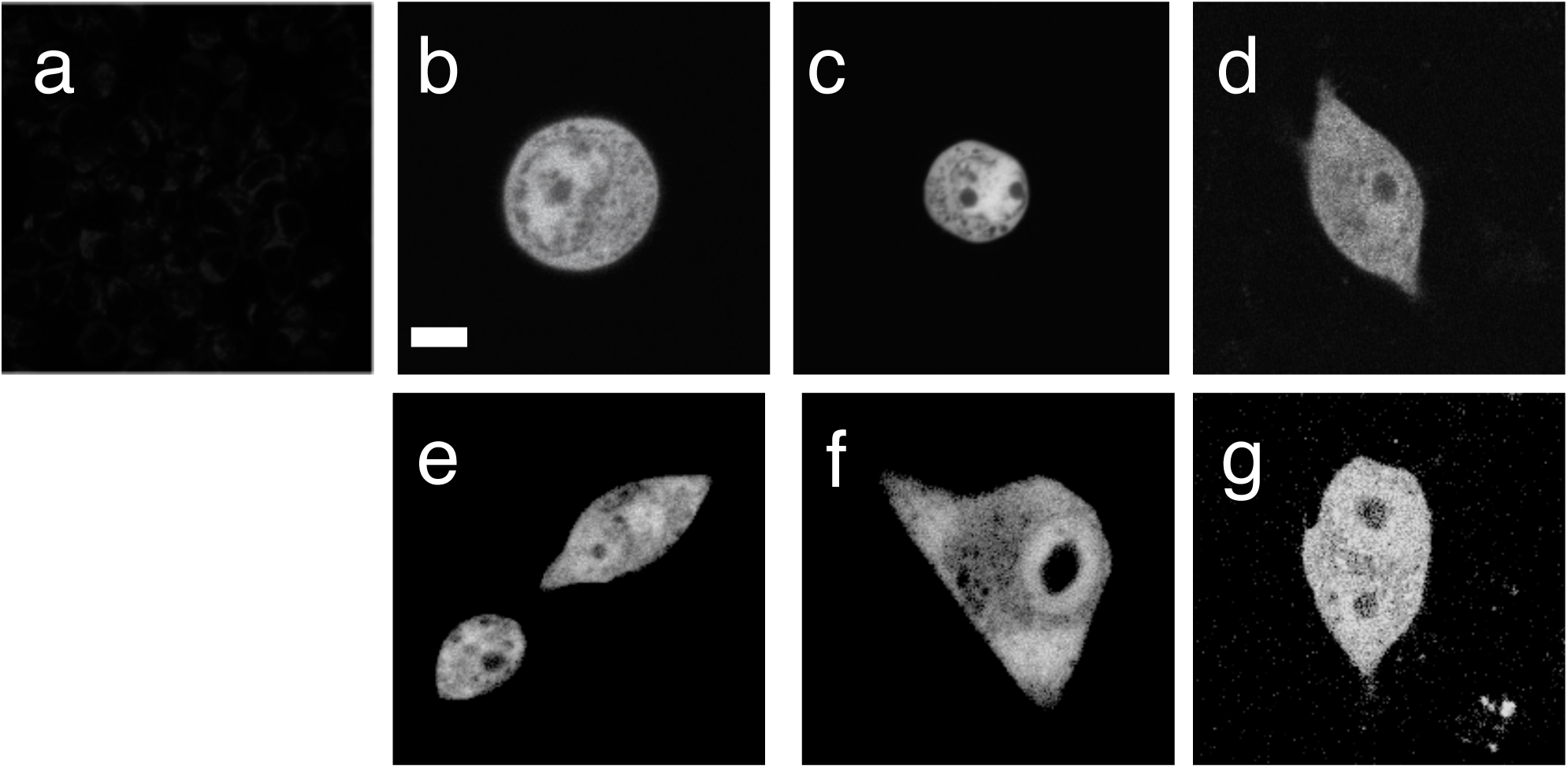
pOV8/Kb complex and chimeric fluorescent protein levels in transfected cell lines. A, F, and G. pOV8/Kb complex on the cell surface was plotted against the fluorescent intensity of the fluorescent protein. Each cell lines indicated left was transfected with a panel of fluorescent proteins indicated above. The surface pOV8/Kb complex was quantified by 25-D1.16 staining 24 hr after transfection for DC2.4 or 48 hr for EL4 and MC57G. The pair of fluorescent histograms from pOV8 containing fluorescent chimeric proteins was compared by color-combined analysis (right-most). (A), The fluorescent histograms of YFP (left-most), pOV8-YFP (left-middle, red), YFP-pOV8 (right-middle, green), and color-combined analysis (right-most). (B), The amount of protein (actin and YFP) in DC2.4 was plotted against the fluorescent intensity of chimeric protein. Each antibody indicated left (anti-actin and anti-GFP clone E4 or RQ2). Antibody against GFP also recognizes YFP, a color variant of GFP. The pair of fluorescent histograms from pOV8 containing fluorescent chimeric proteins were compared by color-combined analysis (right-most). (C), an RT-PCR analysis in DC2.4 comparing transcription of YFP, pOV8-YFP, and YFP-pOV8 (top). Ubiquitin as a loading control (middle). Quantification of RT-PCR (bottom). (D), immunoblot analysis in DC2.4 comparing amounts of YFP, pOV8-YFP, and YFP-pOV8. Immunoblot analysis of YFP, pOV8-YFP, and YFP-pOV8 (top). Anti-Actin as a loading control (middle). Quantification of amounts of YFP, pOV8-YFP, and YFP-pOV8 (bottom). (E), a pulse-chase analysis in DC2.4 comparing stabilities of YFP, pOV8-YFP, YFP-pOV8, and actin under addition of 100 μg/ml of Cycloheximide. Immunoblot analysis comparing amounts of YFP, pOV8-YFP, and YFP-pOV8 (upper-left) and actin as a loading control (lower-left). Quantification of amounts of YFP, pOV8-YFP, and YFP-pOV8 (upper-middle) and actin (lower-middle). The amounts YFP, pOV8-YFP, and YFP-pOV8 were normalized against actin (righ-tmost). (F), The fluorescent histograms of OVA (left-most, red), OVA-YFP (left-middle, red), YFP-OVA (right-middle, green), and color-combined analysis (right-most). (G), The fluorescent histograms of Azamigreen (left-most, red), pOV8-Azamigreen (left-middle, red), Azamigreen-pOV8 (right-middle, green), and color-combined analysis (right-most). (H), Localization of YFP (b), pOV8-YFP (c), YFP-pOV8 (d), Azamigreen (e), pOV8-Azamigreen (f), and Azamigreen-pOV8 (g) in DC2.4. Intracellular localization of fluorescent proteins was detected by fluorescence of each protein. (a) as the control without fluorescent proteins. Bar, 10 μm. Student’s test was used to compare pOV8-YFP sample with YFP-pOV8 sample. Data were shown as mean _±_ S.D. (error bars). NS, not significant, n=3.

**Figure 2.**
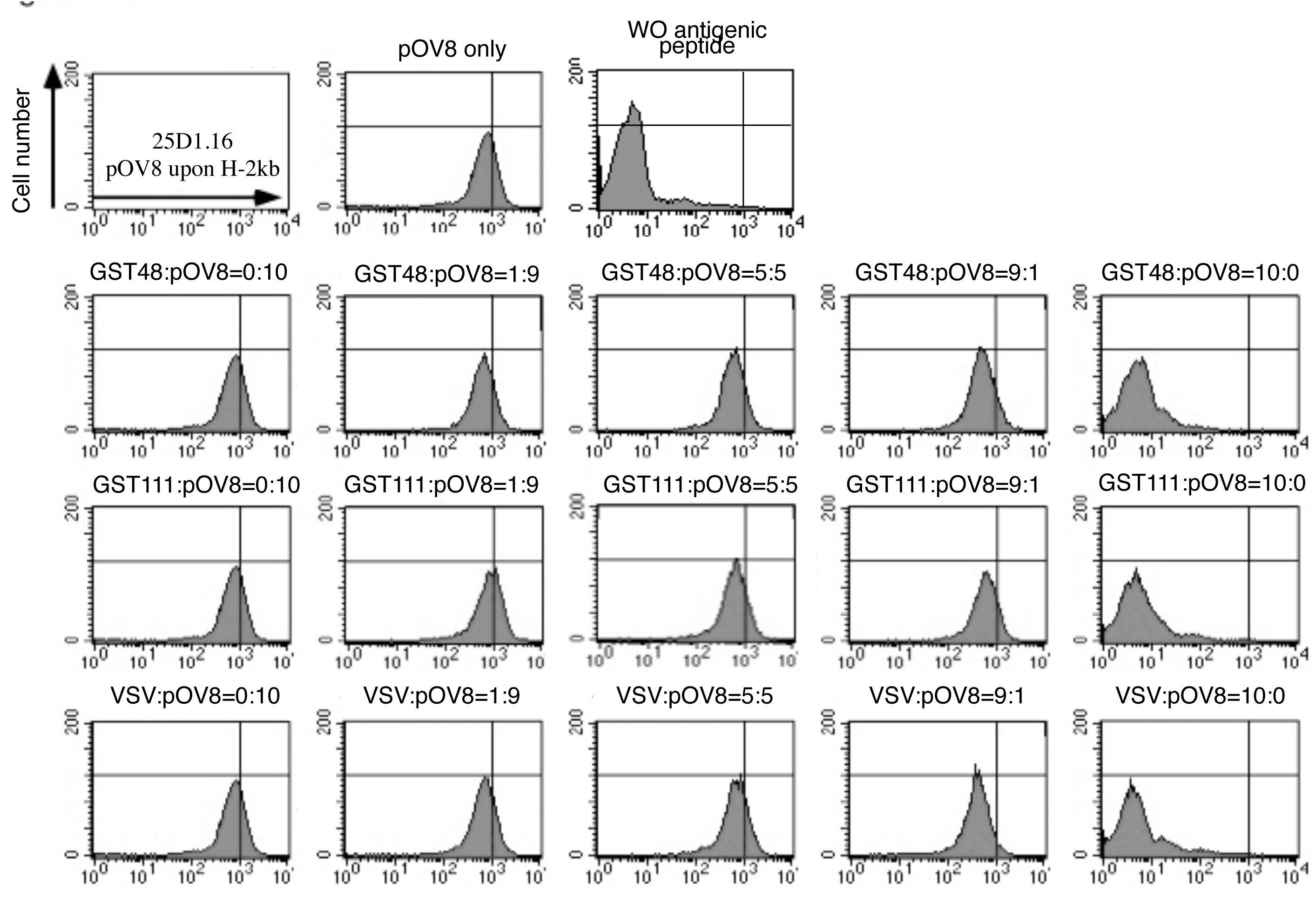
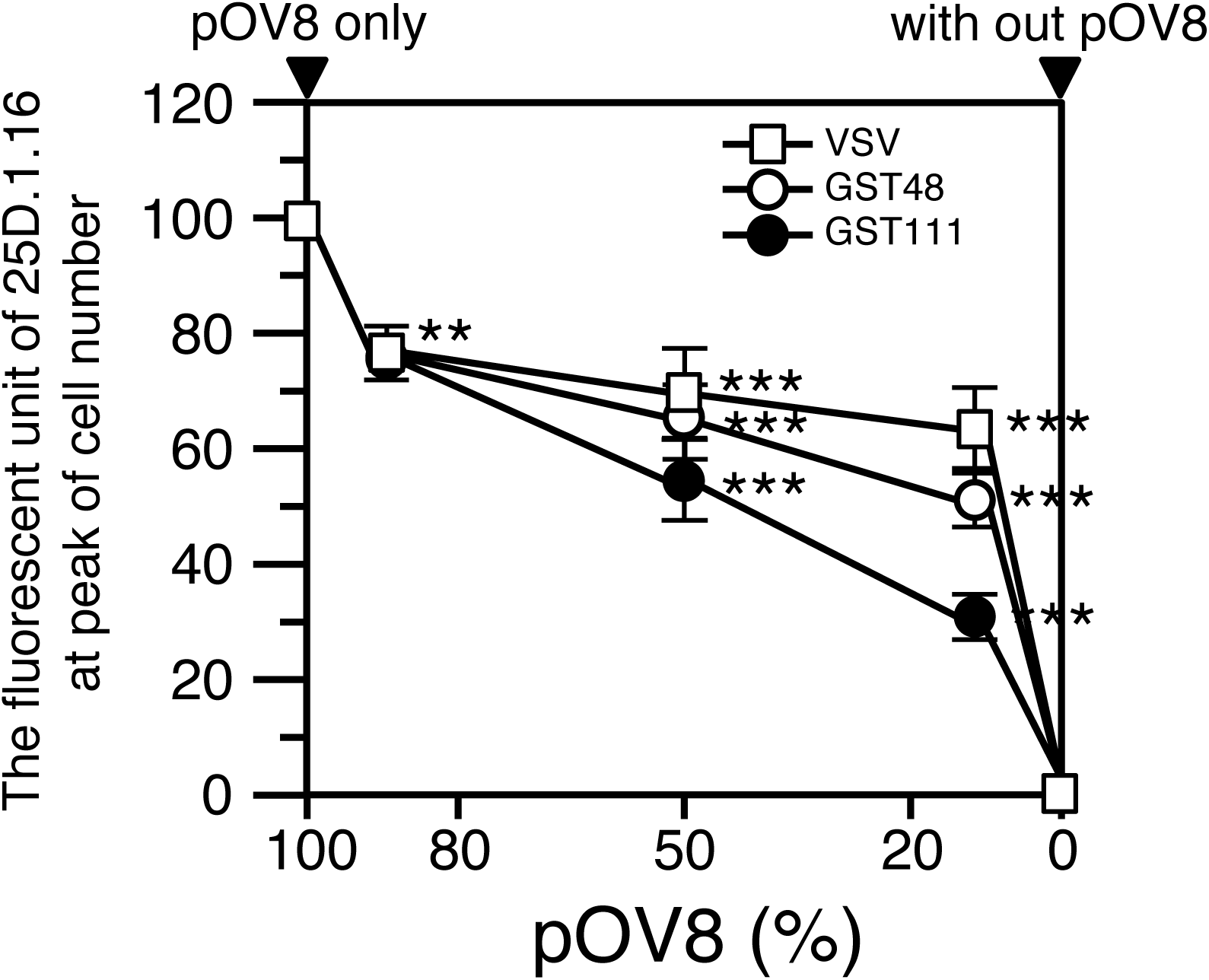
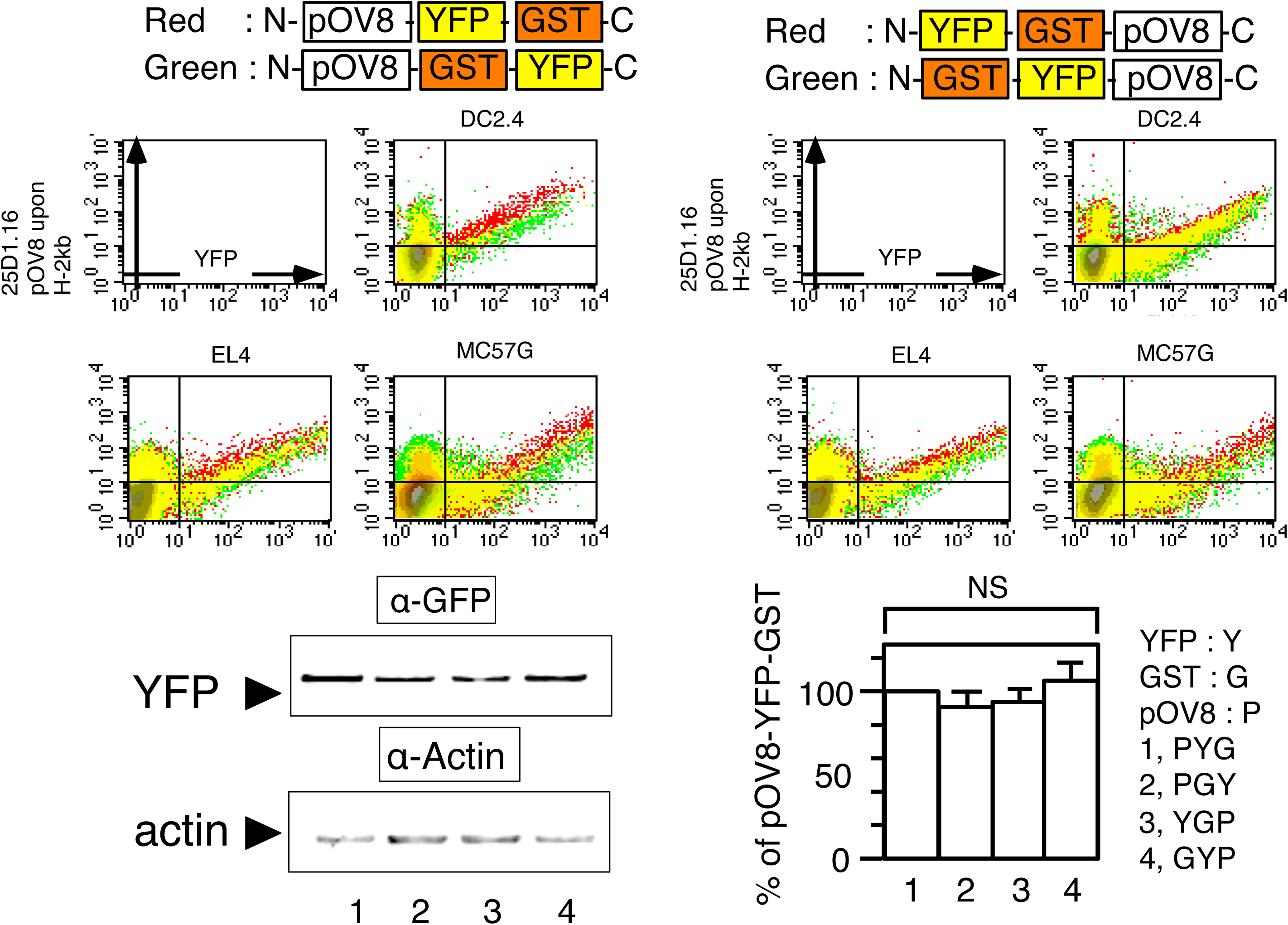
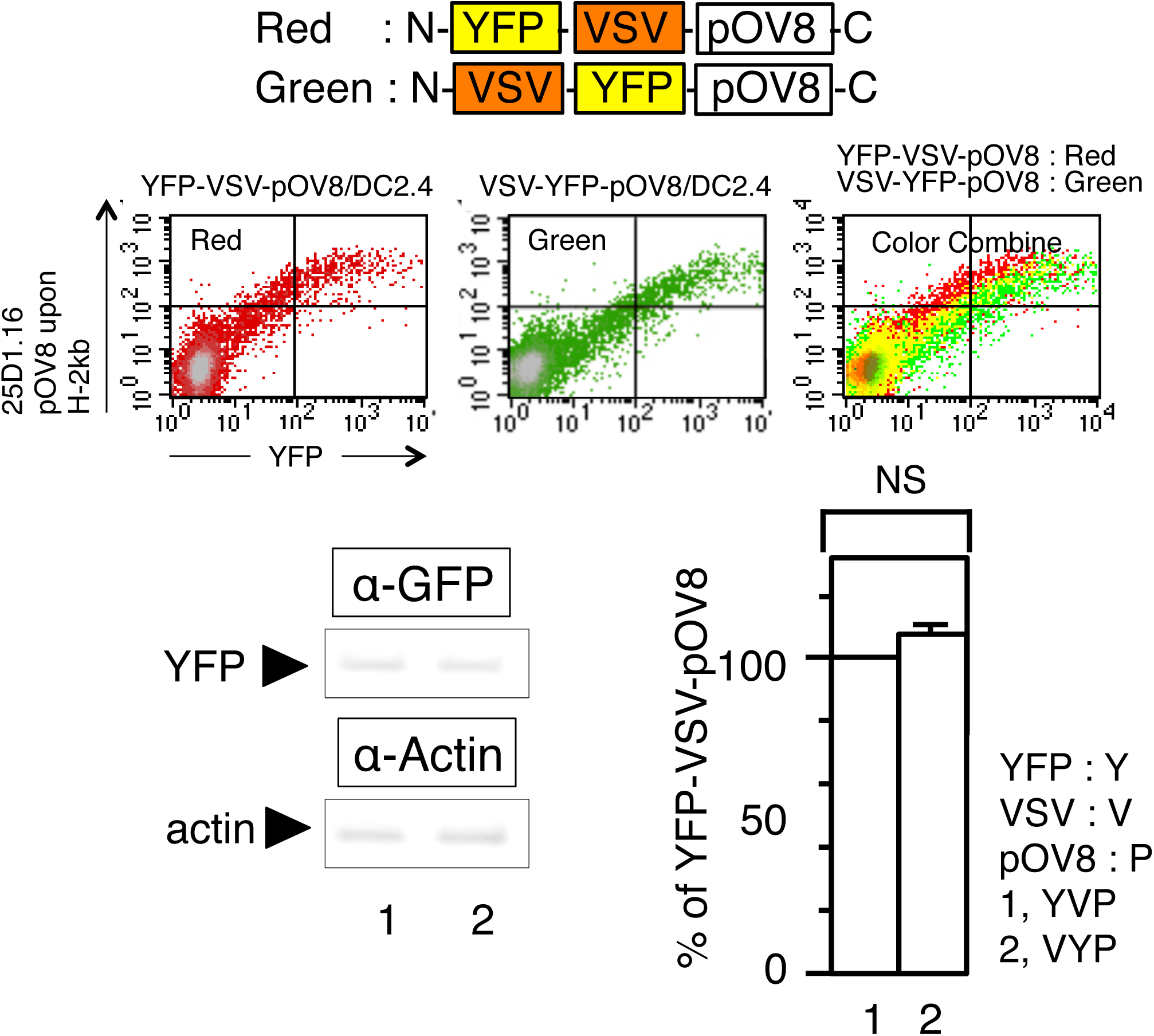
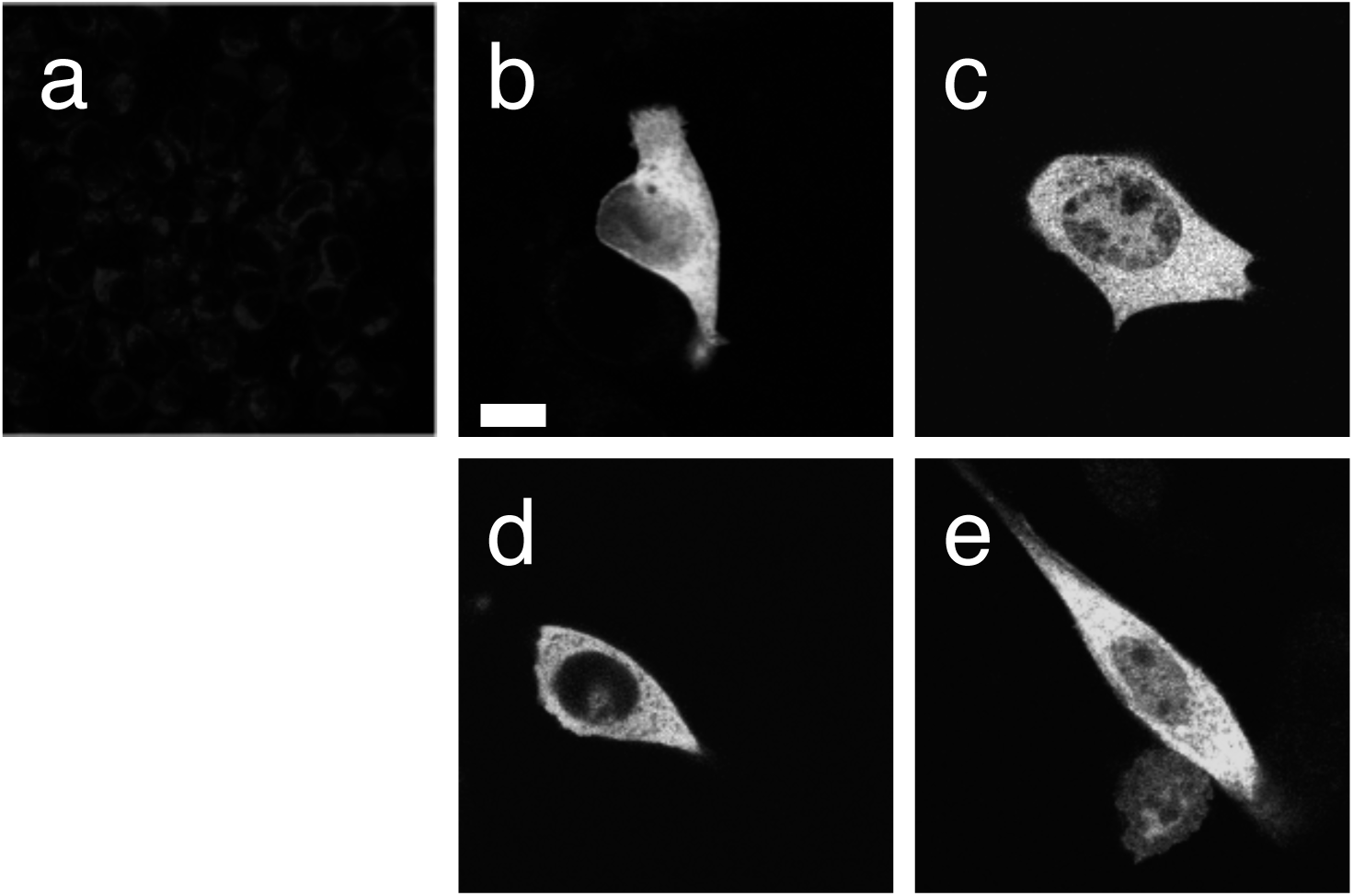
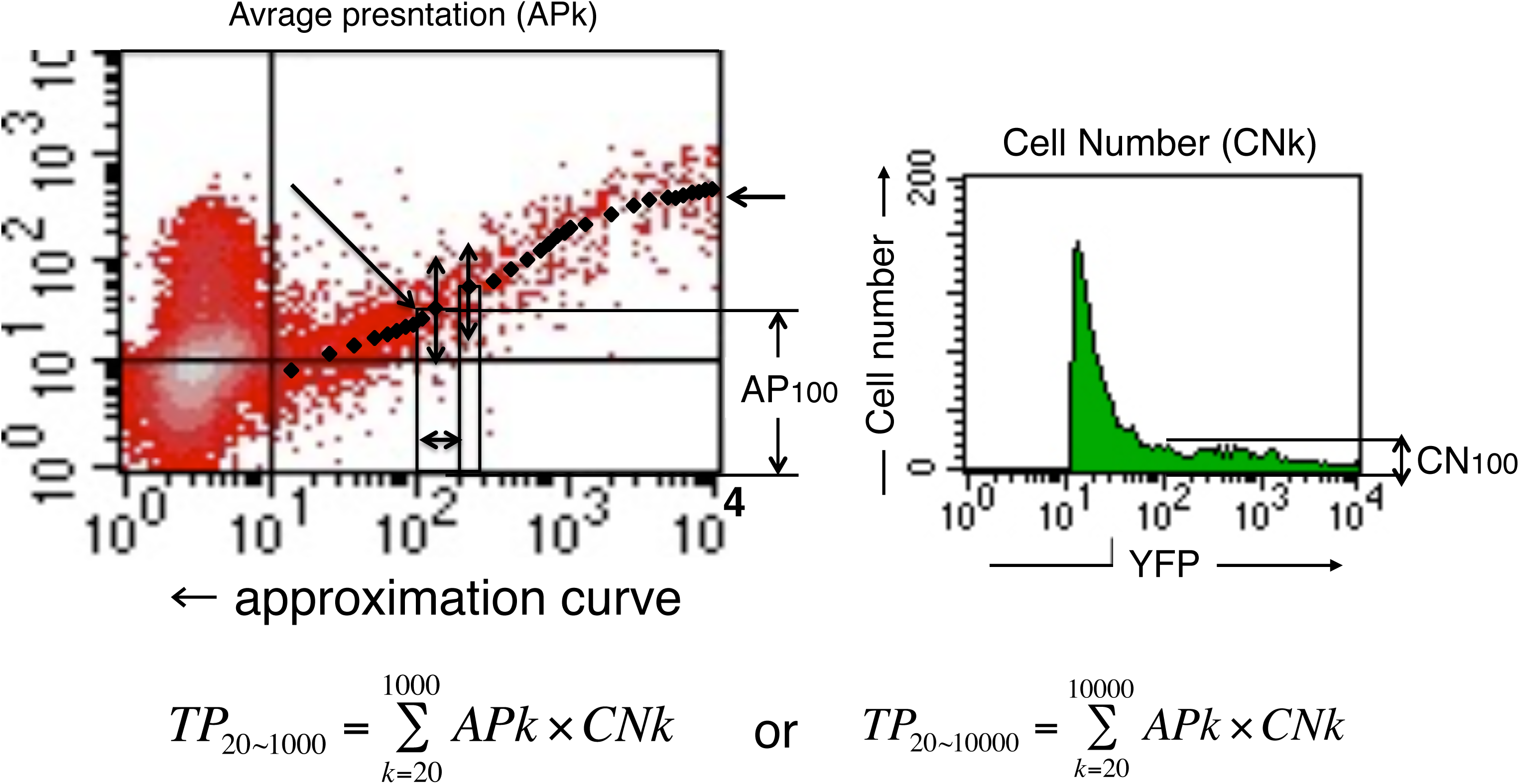
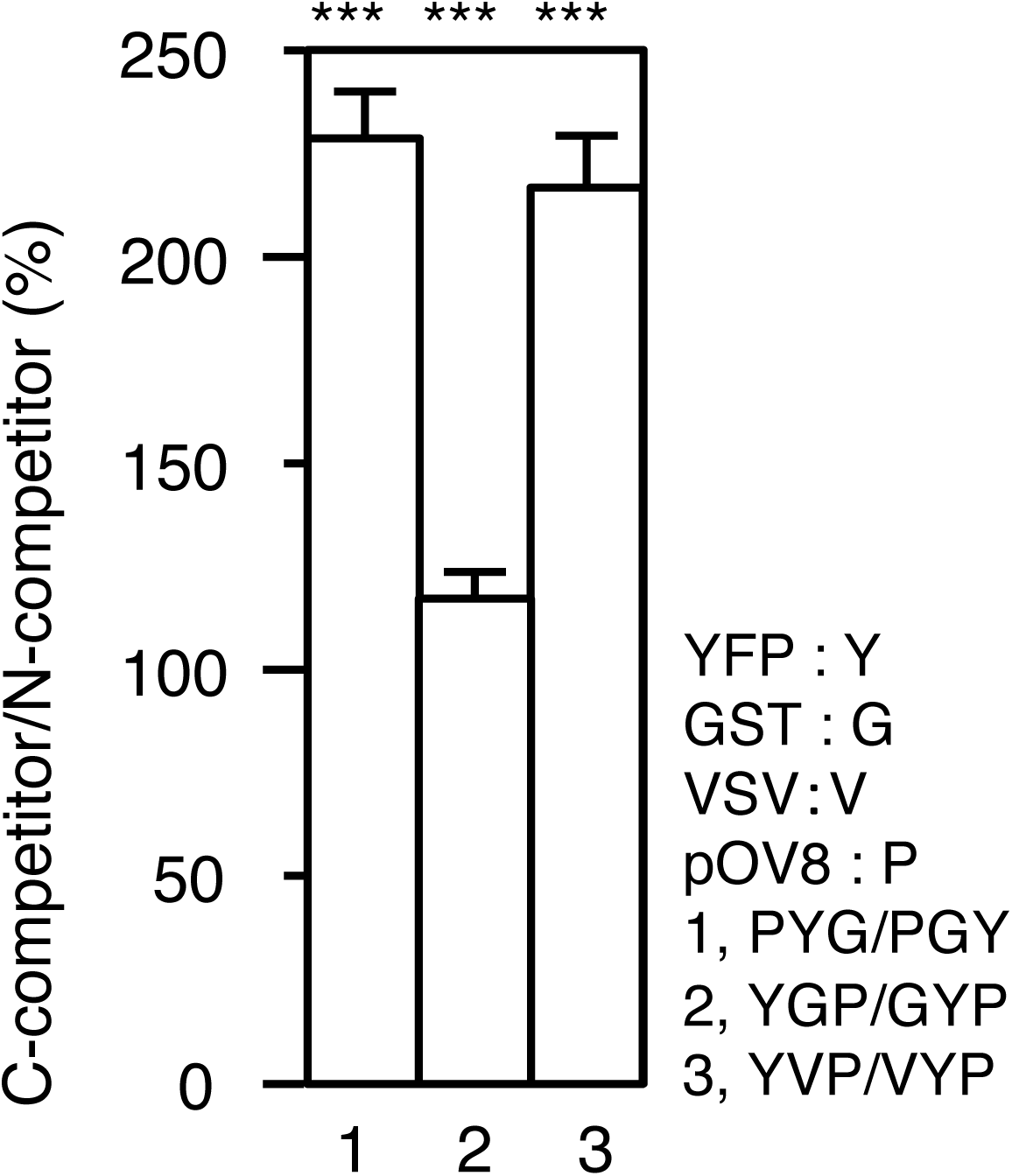
pOV8/Kb complex formations under intramolecular competitions. (A), Flow cytometric analysis of DC2.4 cells pulsed with the constant concentration of peptides (50 μM), in which the decreasing concentrations of pOV8 (50-0 μM) with the increasing concentrations of competitive Ag-peptides (0-50 μM; VSV, GST48, GST111). The relative amounts of each peptide were indicated above each histogram; 50 μM of pOV8 and 0 μM of competitive peptide, 40 μM of pOV8 and 10 μM of competitive peptide, 25 μM of pOV8 and 25 μM of competitive peptide, 10 μM of pOV8 and 40 μM of competitive peptide, and 0 μM of pOV8 and 50 μM of competitive peptide. The surface pOV8/Kb complex was quantified by 25-D1.16. (B), The relative fluorescent unit of horizontal axis-25D.1.16 at the peak of cell numbers from A were plotted against the rate of the pOV8 peptide. (C), The intramolecular competition of the pOV8/Kb complex formations by GST. Each cell lines indicated above was transfected with a pair of fluorescent proteins. Generation of the pOV8/Kb complex from the pair of equivalent transfectants was analyzed by color combine as Fig. 1 (upper-left; red with pOV8-YFP-GST and green with pOV8-GST-YFP. upper-right; red with YFP-GST-pOV8 and green with GST-YFP-pOV8). Immunoblot analysis in DC2.4 comparing amounts of pOVA-YFP-GST, pOVA-GST-YFP, YFP-GST-pOVA, and GST-YFP-pOVA (lower-left). Anti-Actin as a loading control. Quantification of amounts of pOVA-YFP-GST, pOVA-GST-YFP, YFP-GST-pOVA, and GST-YFP-pOVA (lower-right). (D), The intramolecular competition of the pOV8/Kb complex formations by the VSV derived antigenic peptides. DC2.4 was transfected with a pair of fluorescent proteins. Generation of the pOV8/Kb complex from the pair of equivalent transfectants was analyzed by color combine as Fig. 1 (upper-left; red with YFP-VSV-pOV8, upper-middle; green with VSV-YFP-pOV8, and upper-left; color combine). Immunoblot analysis comparing amounts of YFP-VSV-pOV8 and VSV-YFP-pOV8 (bottom-left). Anti-Actin as a loading control. Quantification of amounts of YFP-VSV-pOV8 and VSV-YFP-pOV8 (bottom-right). (E), Localization of pOVA-YFP-GST (b), pOVA-GST-YFP (c), YFP-GST-pOVA (d), and GST-YFP-pOVA (e) in DC2.4. Intracellular localization of fluorescent protein was detected by fluorescence of YFP. (a) as the control without fluorescent proteins. Bar, 10 μm. (F), Quantification of the amounts of pOV8/Kb complexes. Amounts of the pOV8/Kb complex of each experiment were worked out as Material and Methods. (G), Quantification of decreased presentation of the pOV8/Kb complex of DC2.4 by competition with GST derived peptides (TP_20∼10000_) and the VSV antigenic peptide(TP_20∼10000_) shown in C and D by DC2.4. Student’s test was used to compare the quantified results. Data were shown as mean _±_ S.D. (error bars). ***, p<0.01, **,p<0.1, and NS, not significant. n=3.

**Figure 3.**
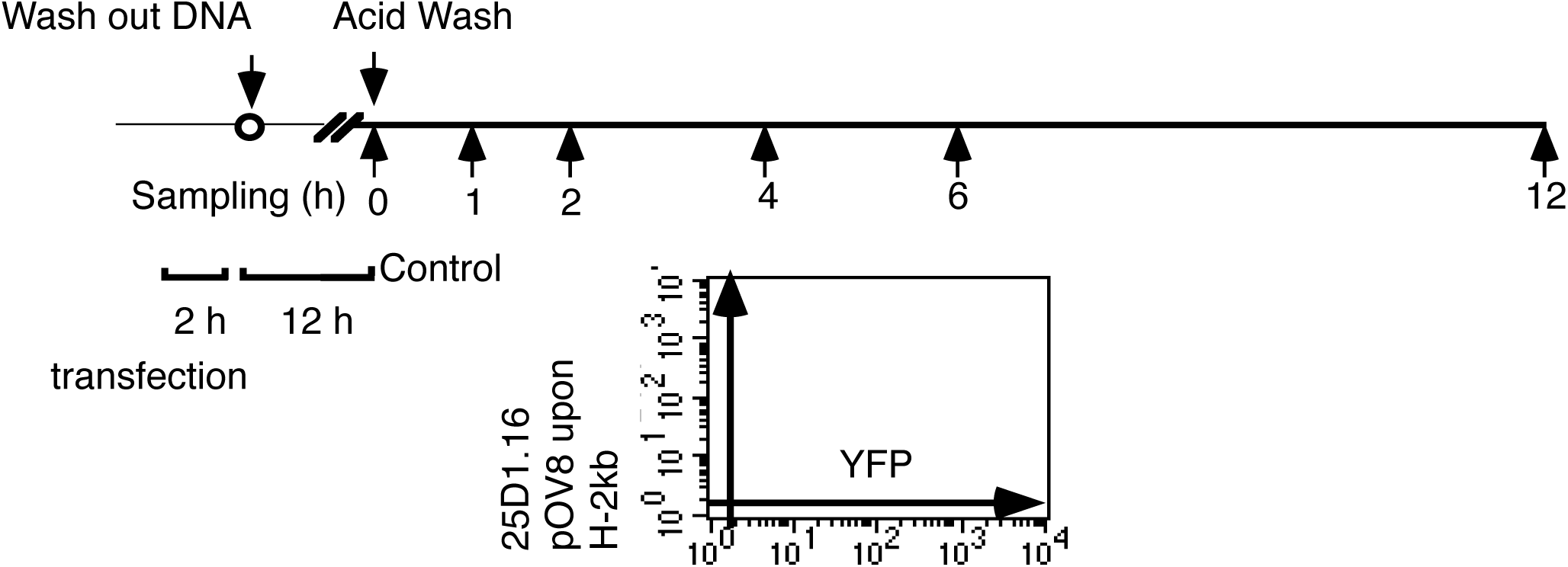
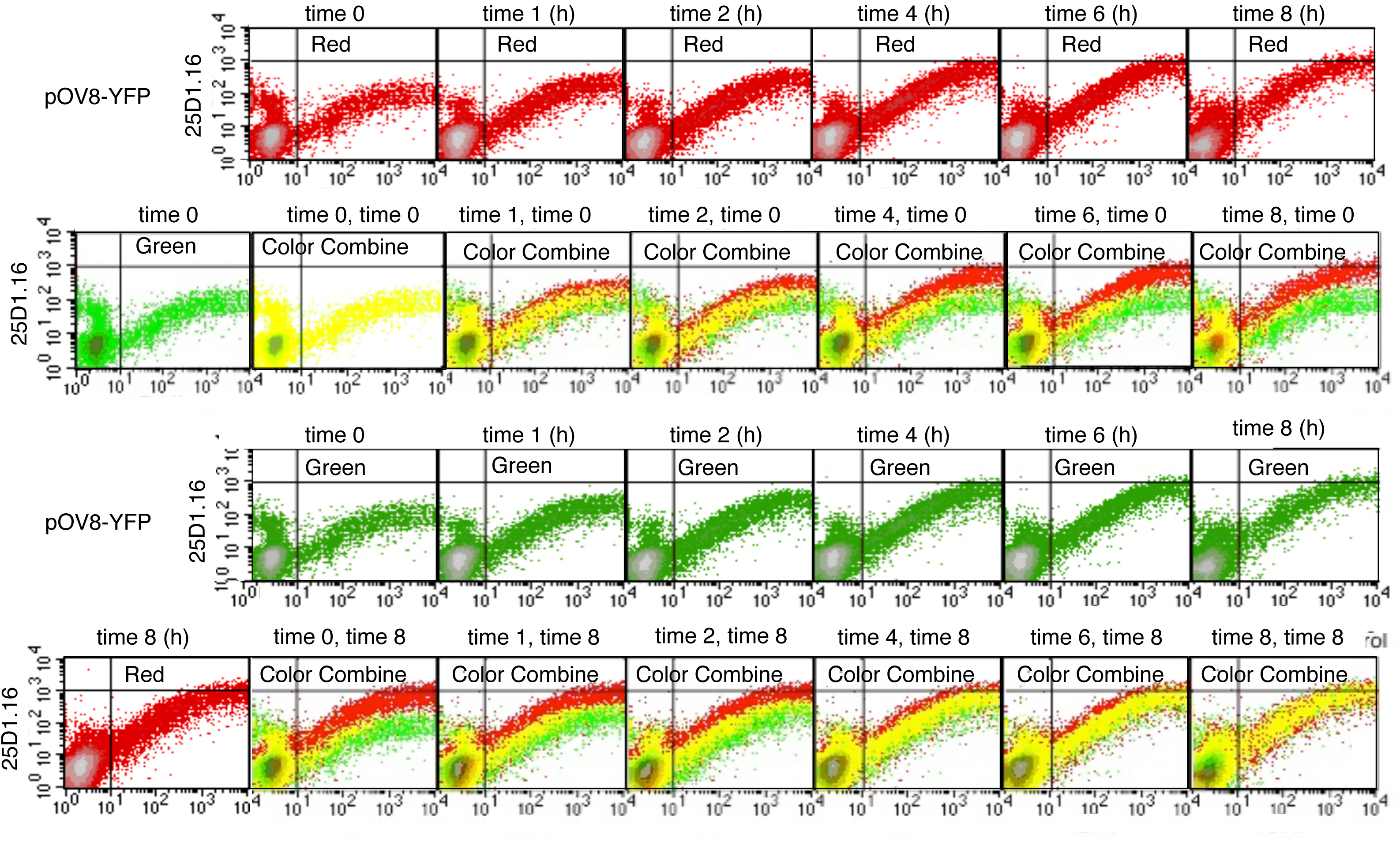
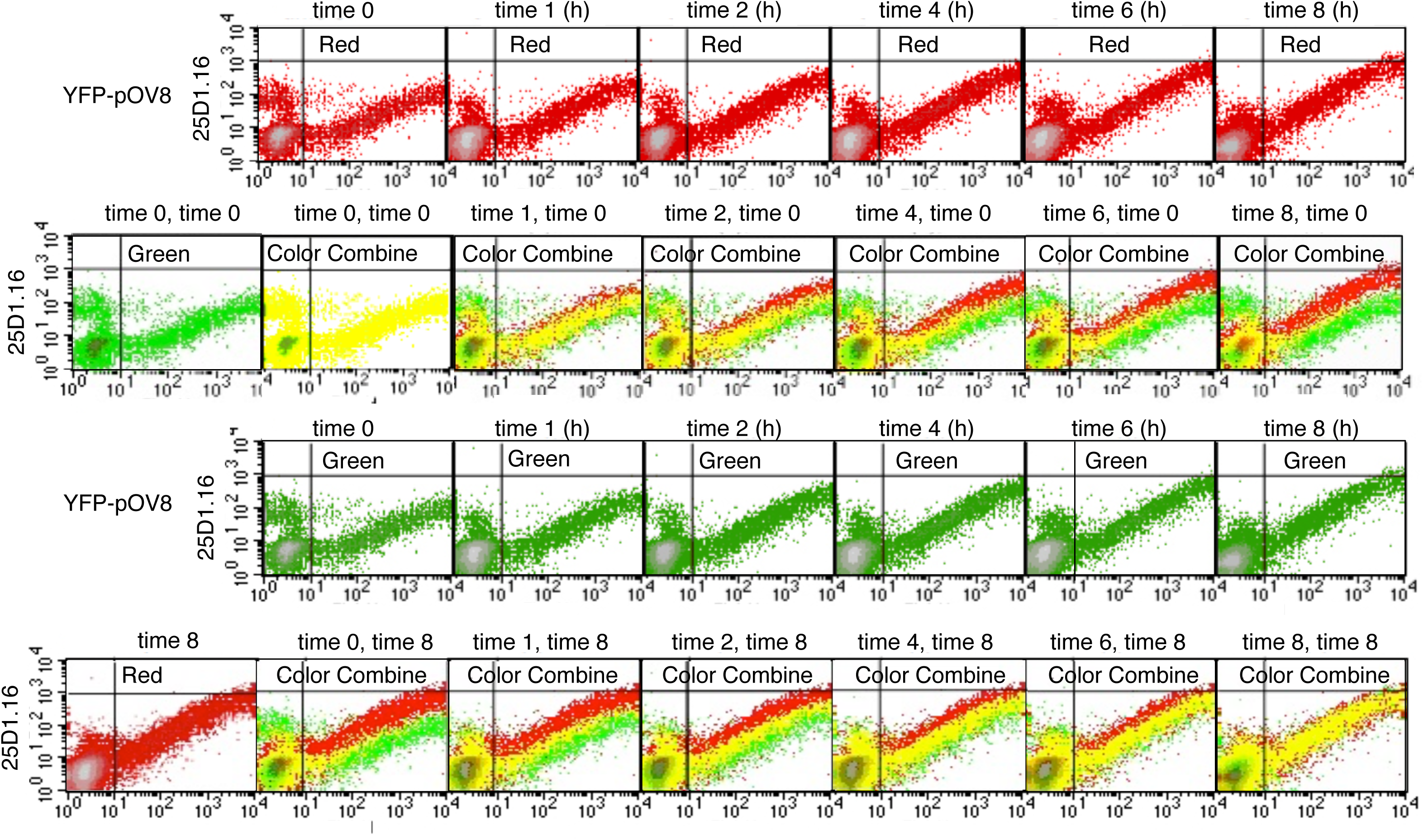
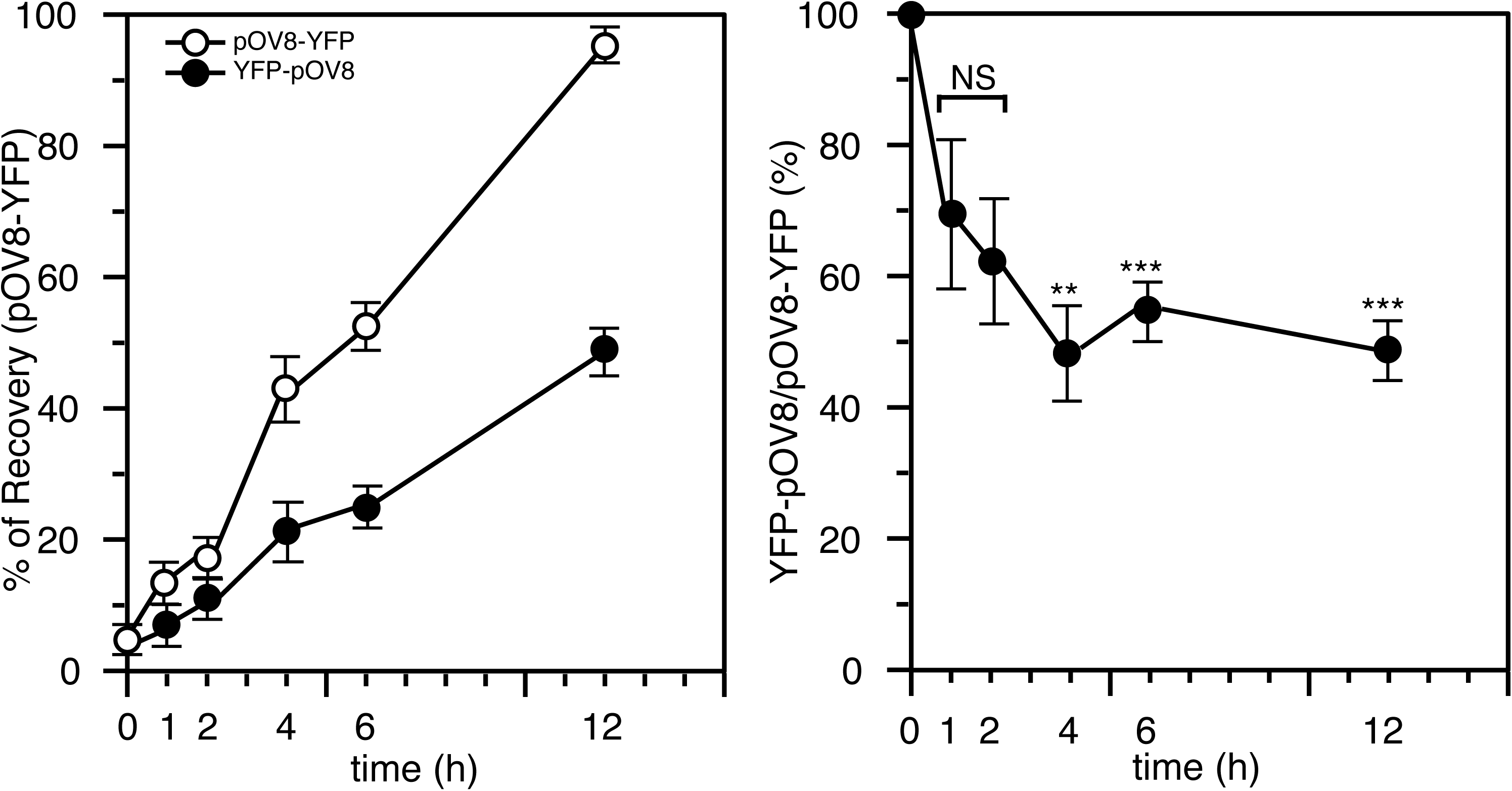
Kinetics of generation of the surface pOV8/Kb complex. (A), DC2.4 cells were acid washed after 12 h of transfection. Cells without acid wash were sampled as a control. Cells were sampled at each time point after the acid wash. B and C, Generation of pOV8/Kb complex in DC2.4 cells transfected with pOV8-YFP (B) and YFP-pOV8 (C). Upper lane; fluorescent histograms for the generation of the pOV8/Kb complex in each sample (red). Second lane; fluorescent histograms of each sample (red, first lane) were compared with the sample of time 0 (green, left) by a color combine. Third lane; fluorescent histograms for the generation of the pOV8/Kb complex in each sample (green). Lower lane; fluorescent histograms of each sample (green, third lane) was compared with the sample of control (red, left) by a color combine. (D), Surface pOV8/Kb complex levels were quantified by the average expression for each sample and indicated as a percent of the control of pOV8-YFP as Fig. 2F. Note that, though quantitative analyses show equivalent linear kinetics from three independent experiments, results shows significant difference among each experiment, because of cellar or staining conditions never strictly agrees with each other. A typical the pair of a result was indicated by the deviation of each quantification to show the linear kinetics. Student’s test was used to compare the pOVA-YFP sample with corresponding YFP-pOVA sample. Data were shown as mean _±_ S.D. (error bars). ***, p<0.01, **,p<0.1, and NS, not significant. n=3.

**Figure 4.**
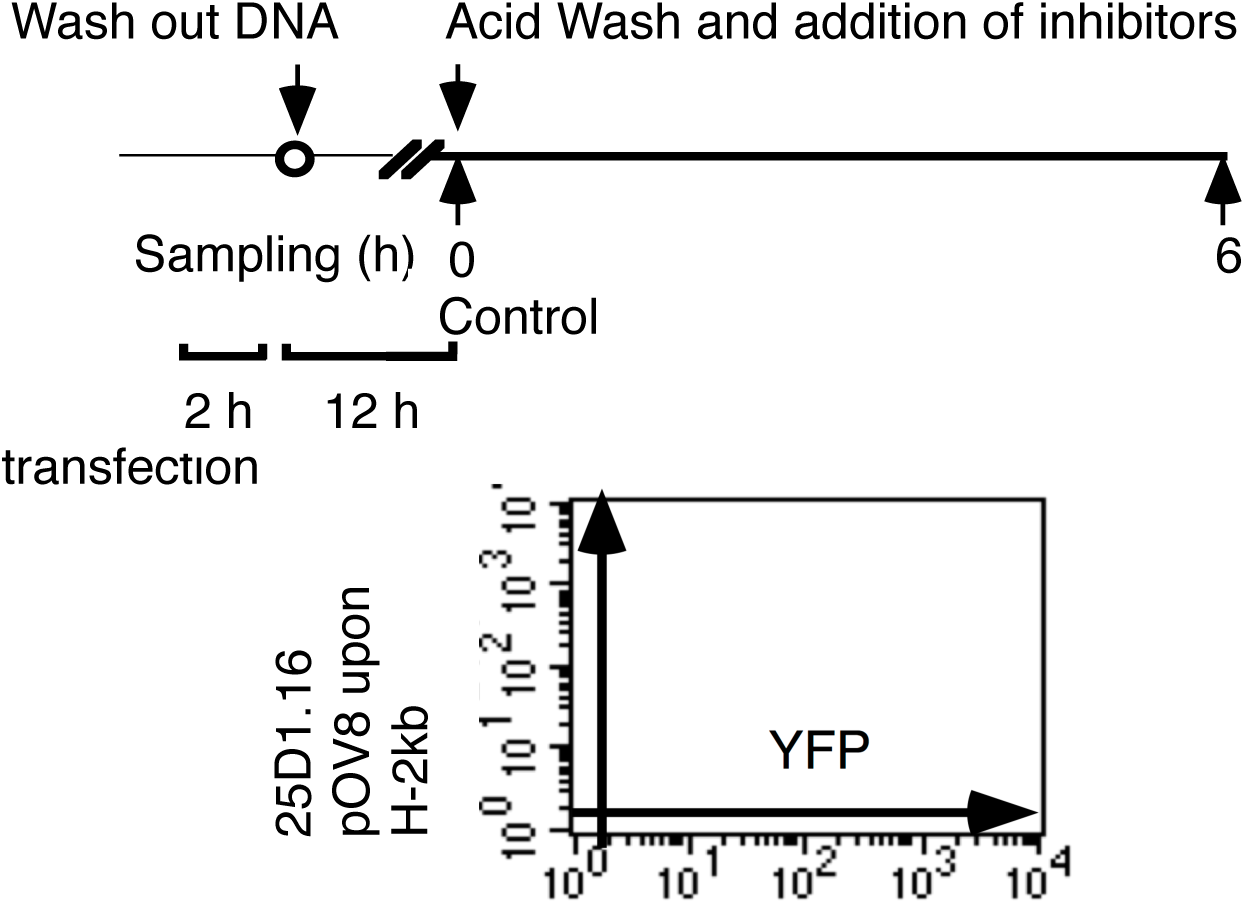
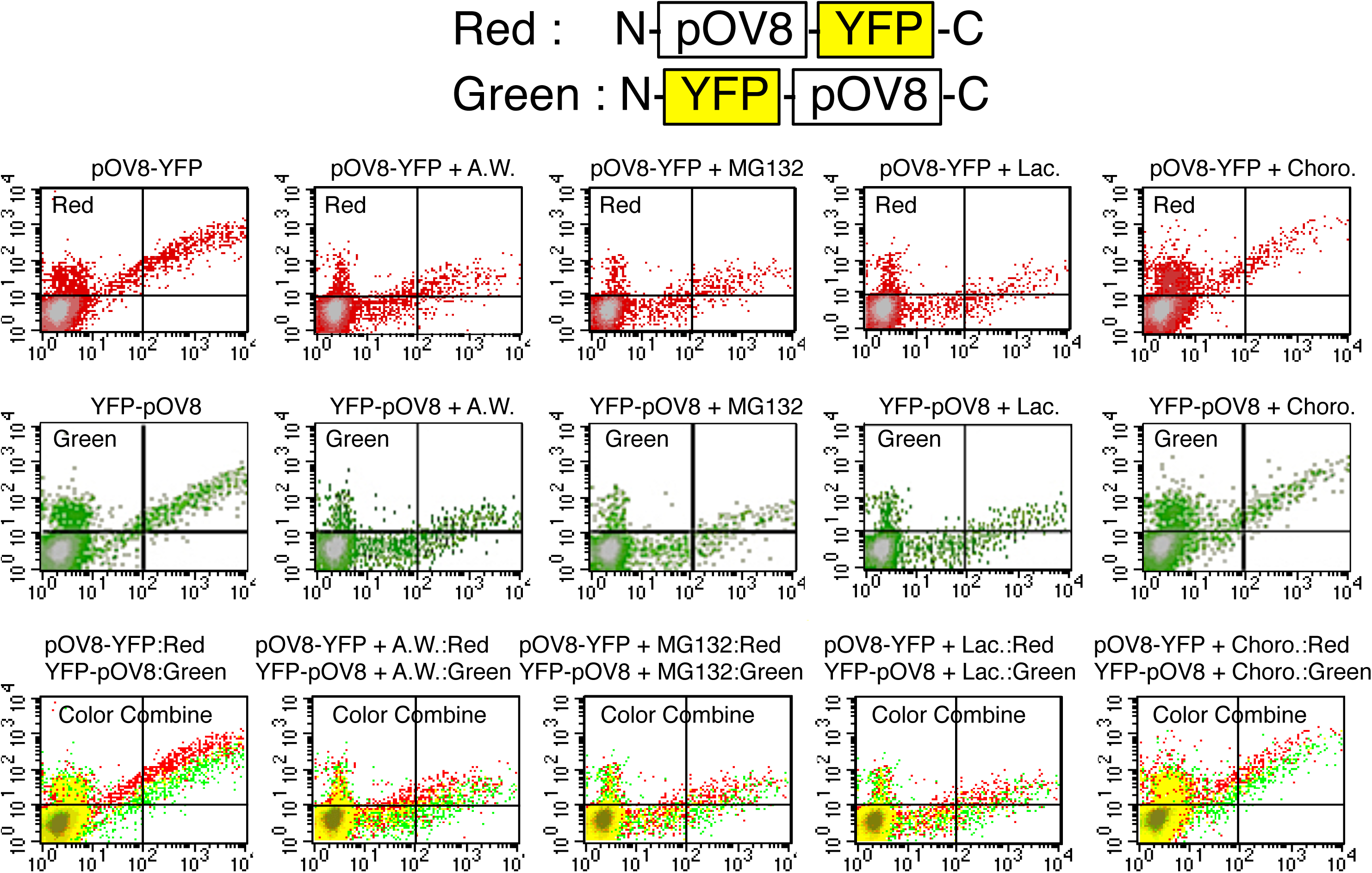
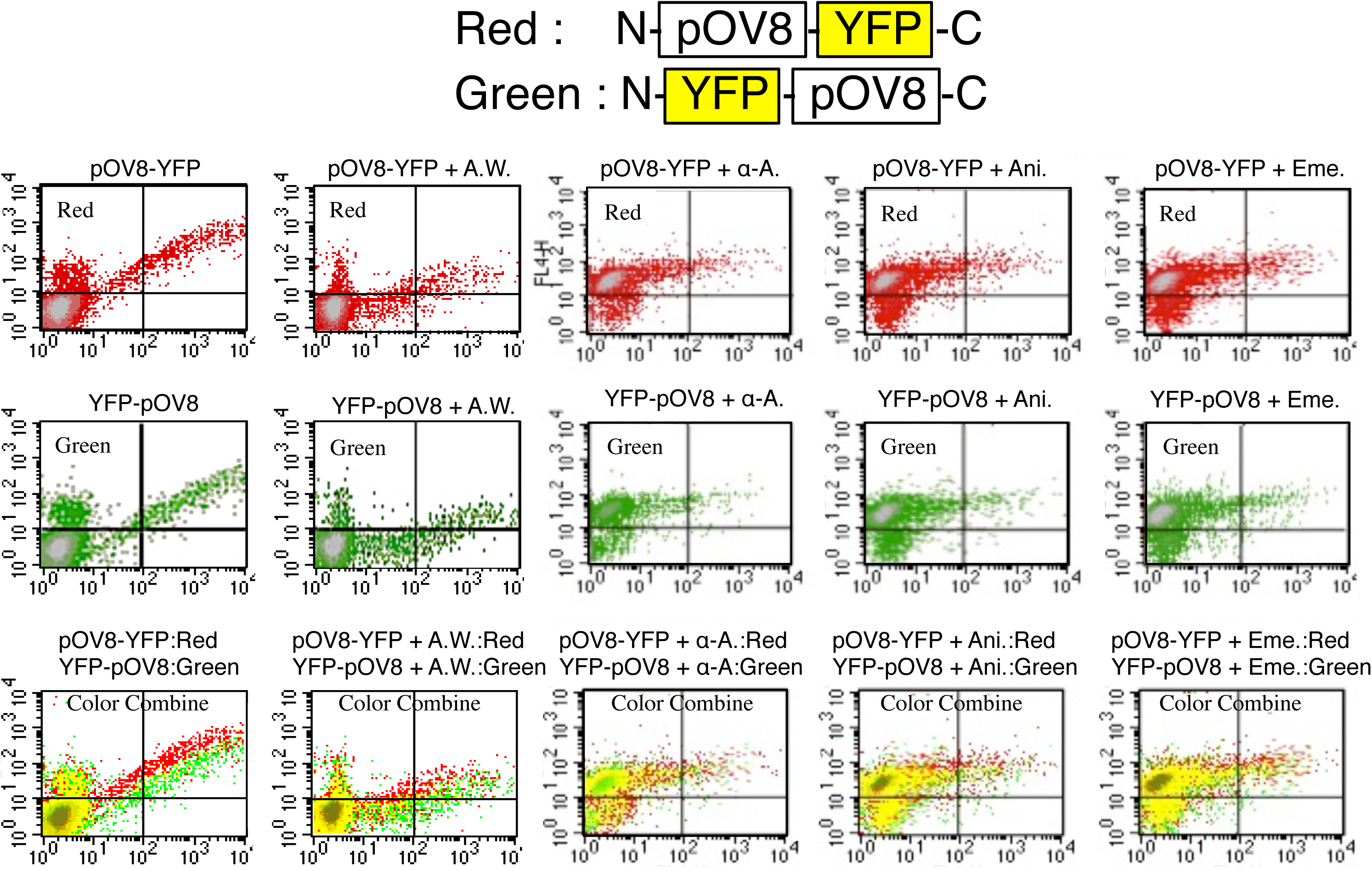
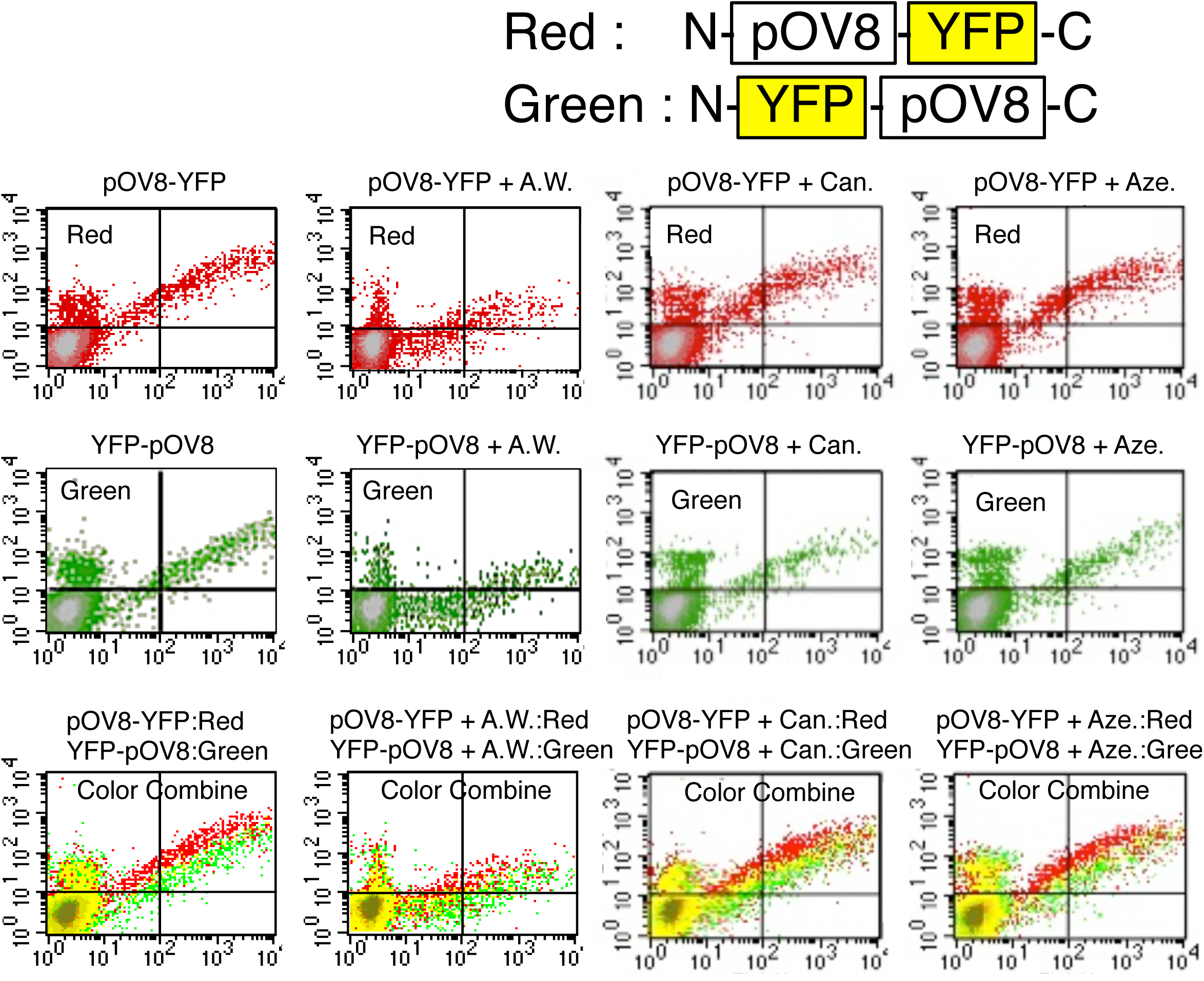
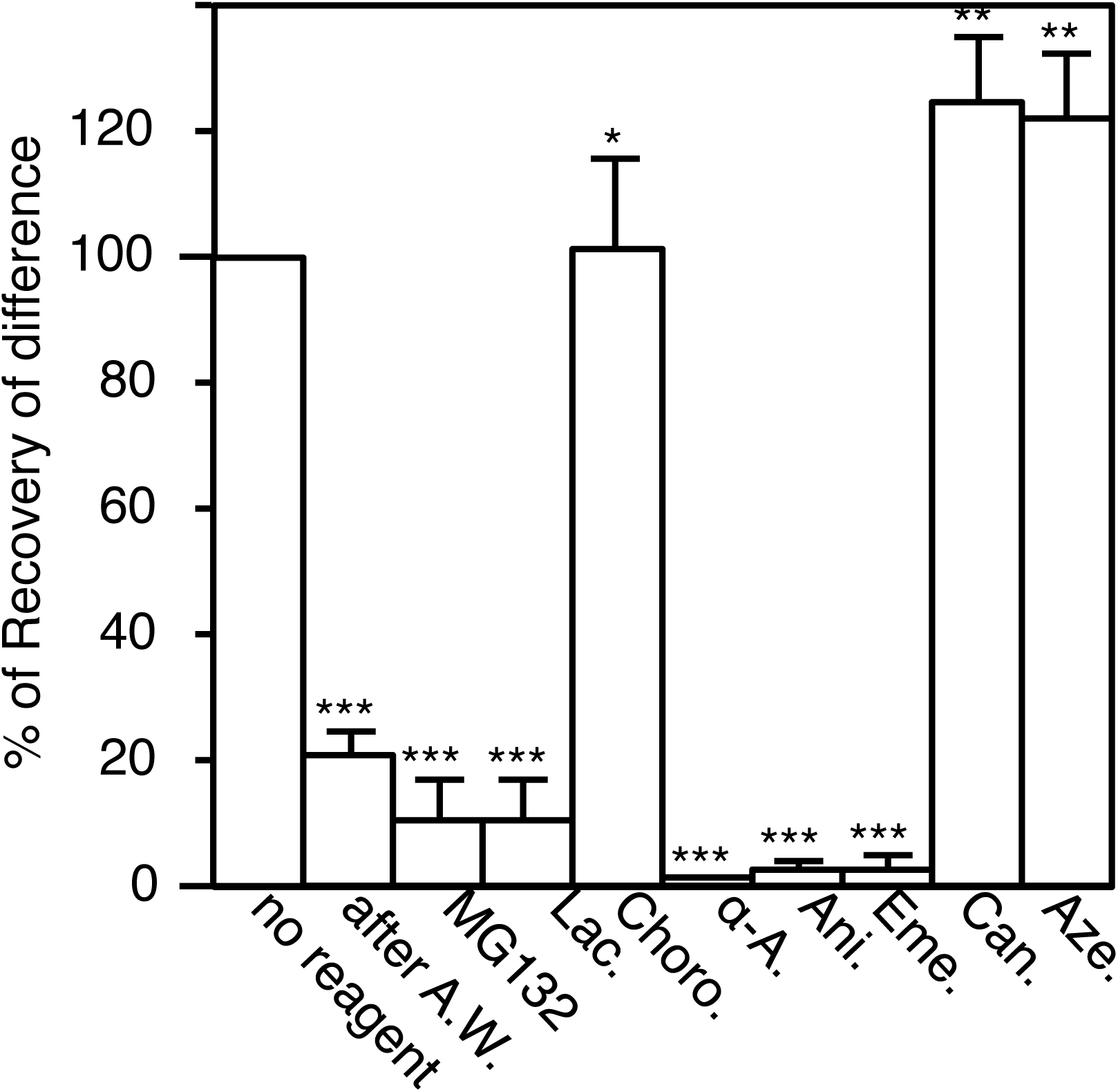
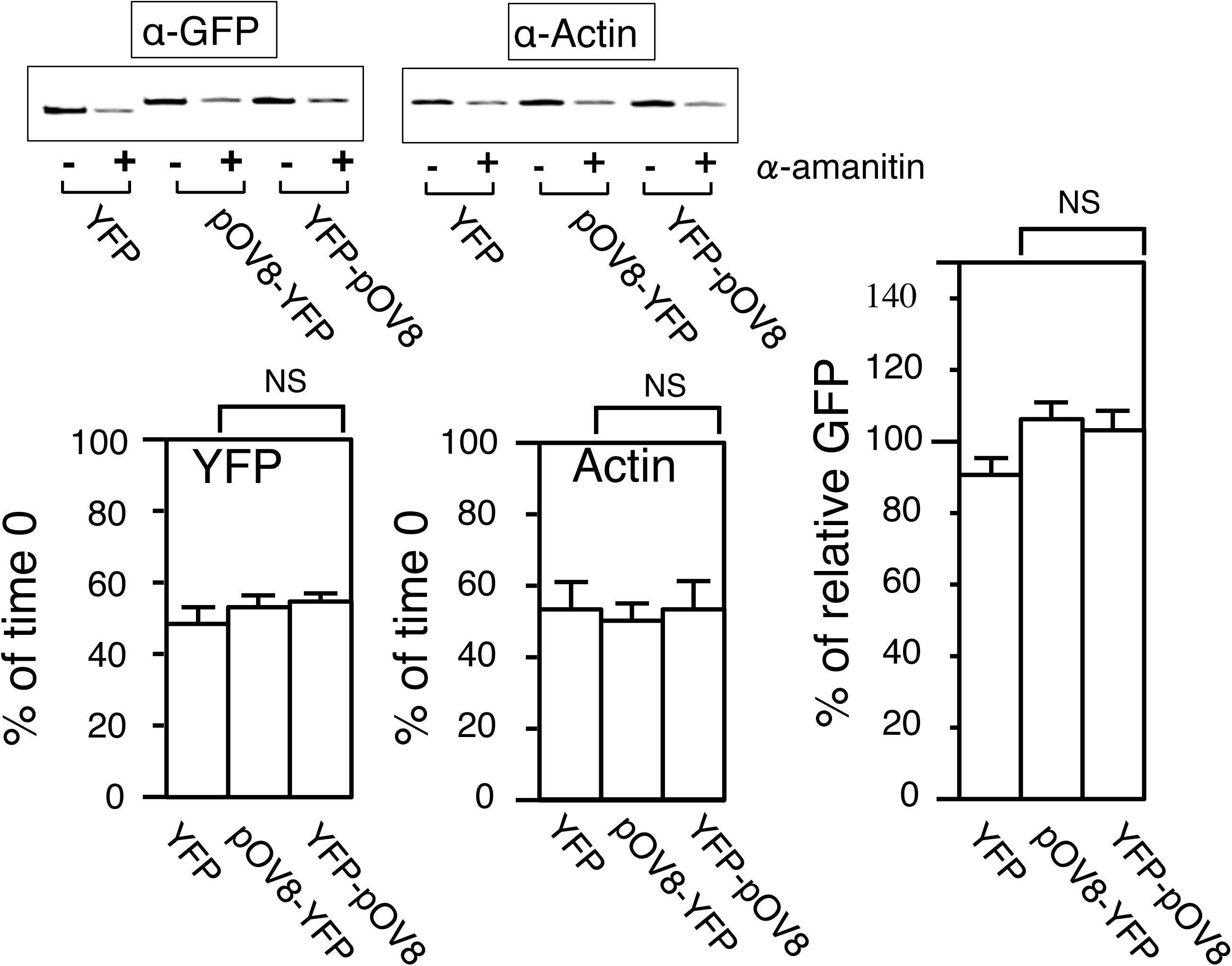
Effects on pOV8/Kb complex generation by different inhibitors altering protein degradation or synthesis. (A), DC2.4 cells were acid washed after 12 h of transfection as in Fig. 3. Cells were cultured for 6 h in normal medium with or without inhibitors. pOV8/Kb complex generation was analyzed as Fig.1. (B), Effects of inhibitors for protein degradation on the production of the pOV8/Kb complex. DC2.4 cells transfected with pOV8-YFP or YFP-pOV8 were cultured with or without 1 μM of MG132, 0.2 μM of lactacystin, or 100 μM of chloroquine. Upper lane; fluorescent histograms of cells transfected with pOV8-YFP. Middle lane; fluorescent histograms of cells transfected with YFP-pOV8. Lower lane; The color combine analyses of above two histograms coloring pOV8-YFP transfected cells as red and YFP-pOV8 transfected cells as green. (C), Effects of inhibitors for protein synthesis. Cells as in B were cultured with or without 10 μg/ml of _α_-amanitin, 1 μg/ml anisomycin, 1 μg/ml emetine. Each fluorescent histograms were analyzed as in B. (D), Effects of inhibitors for protein folding. Cells as in B were cultured with or without 15 mM of canavanine, or 15 mM of azetidine acid. Each fluorescent histograms were analyzed as in B. (E), Quantification for effects reagents upon presentation of the pOV8/Kb complex in B, C, and D as Fig. 2F. (F), immunoblot analysis comparing amounts of YFP, pOVA-YFP, and YFP- pOVA, with or without 10 μg/ml of _α_-amanitin for 6 h (left-top) and quantification of the immunoblot by _α_-GFP (left-bottom). Anti-Actin as a loading control (middle-top) and quantification of the immunoblot by _α_-actin (middle-bottom). The amounts YFP, pOV8-YFP, and YFP-pOV8 were normalized against actin (right-most). Student’s test was used to compare corresponding pOVA-YFP and YFP-pOVA. Data were shown as mean _±_ S.D. (error bars). ***, p<0.01, **,p<0.1, and NS, not significant. n=3.

**Figure 5.**
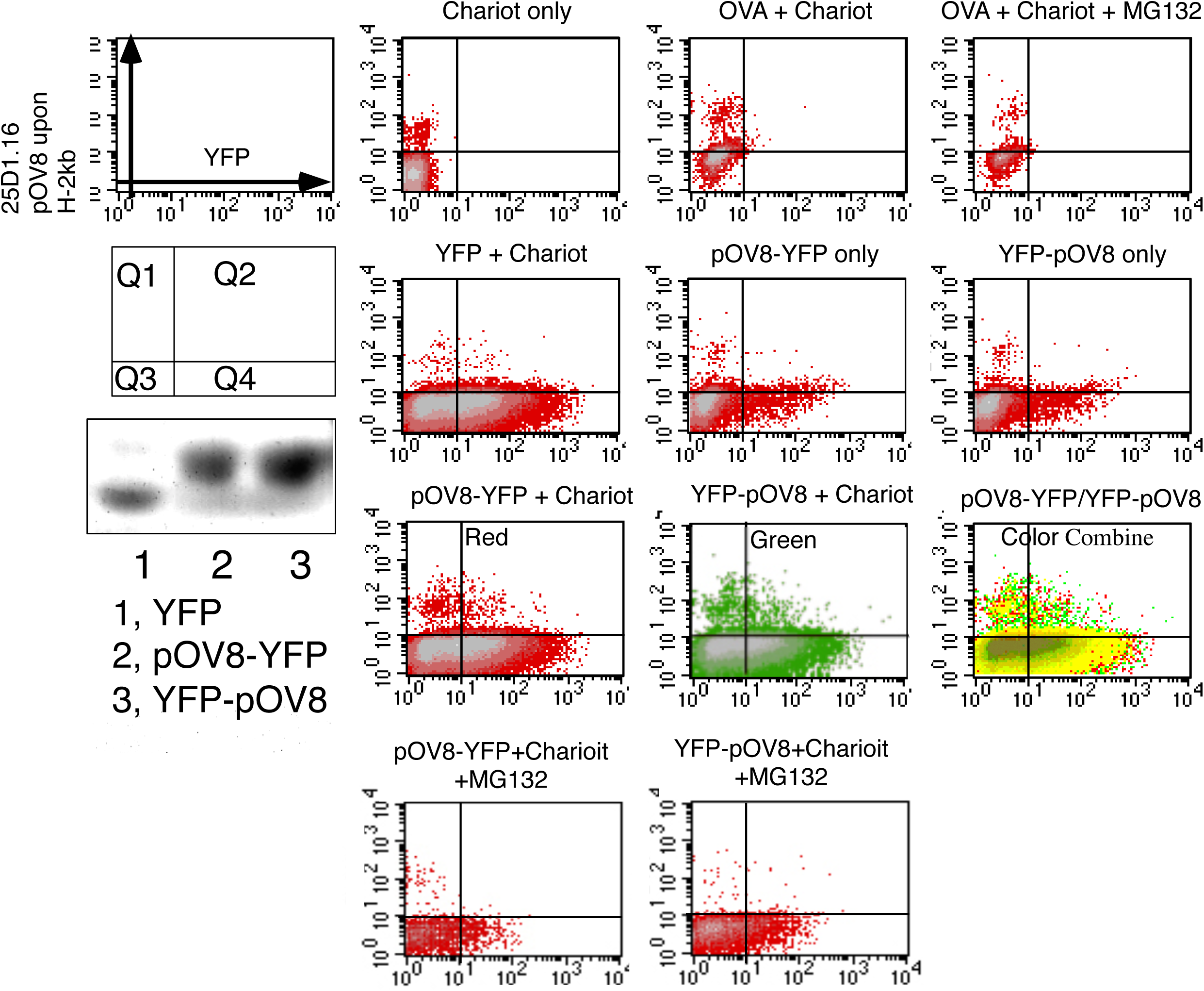
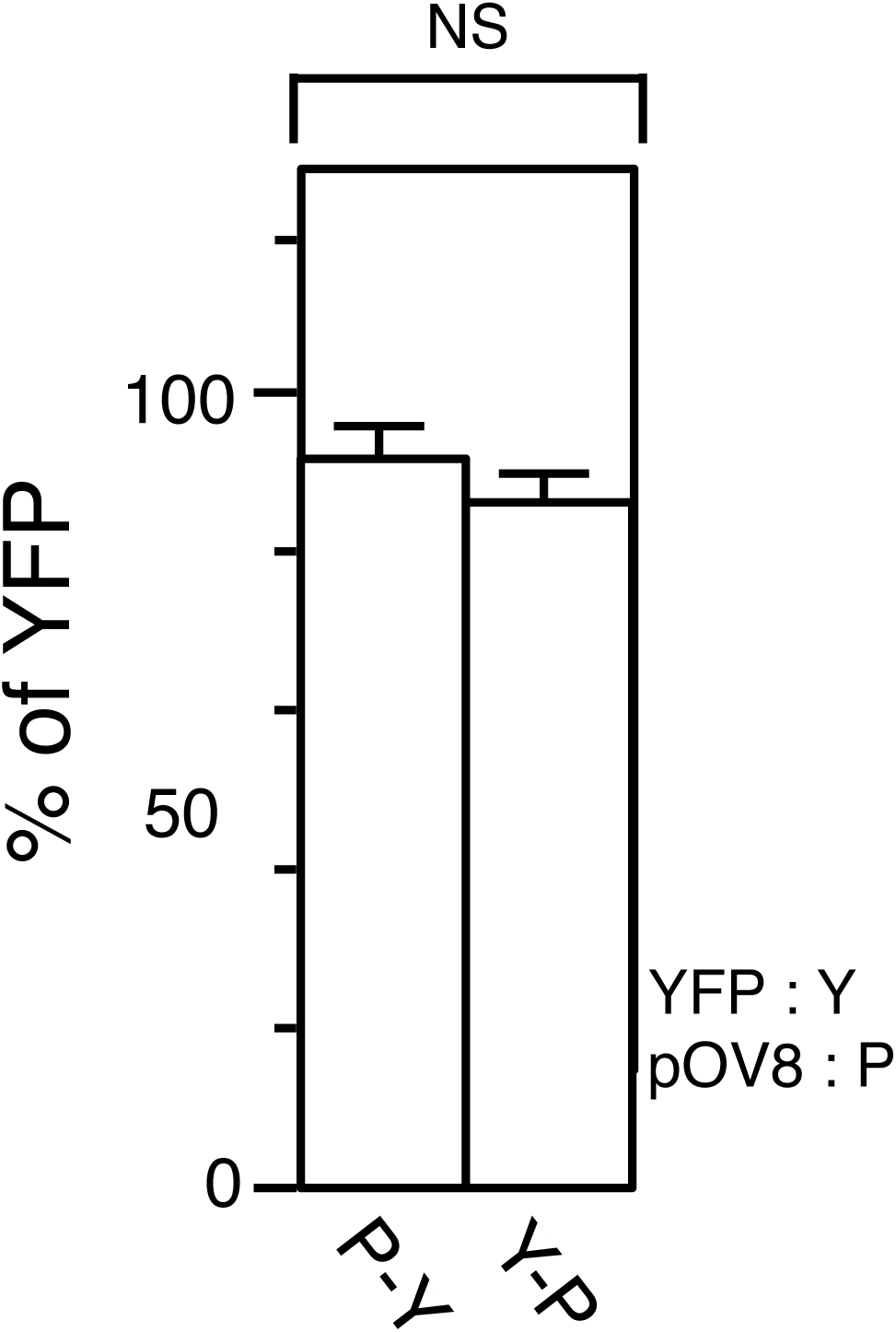
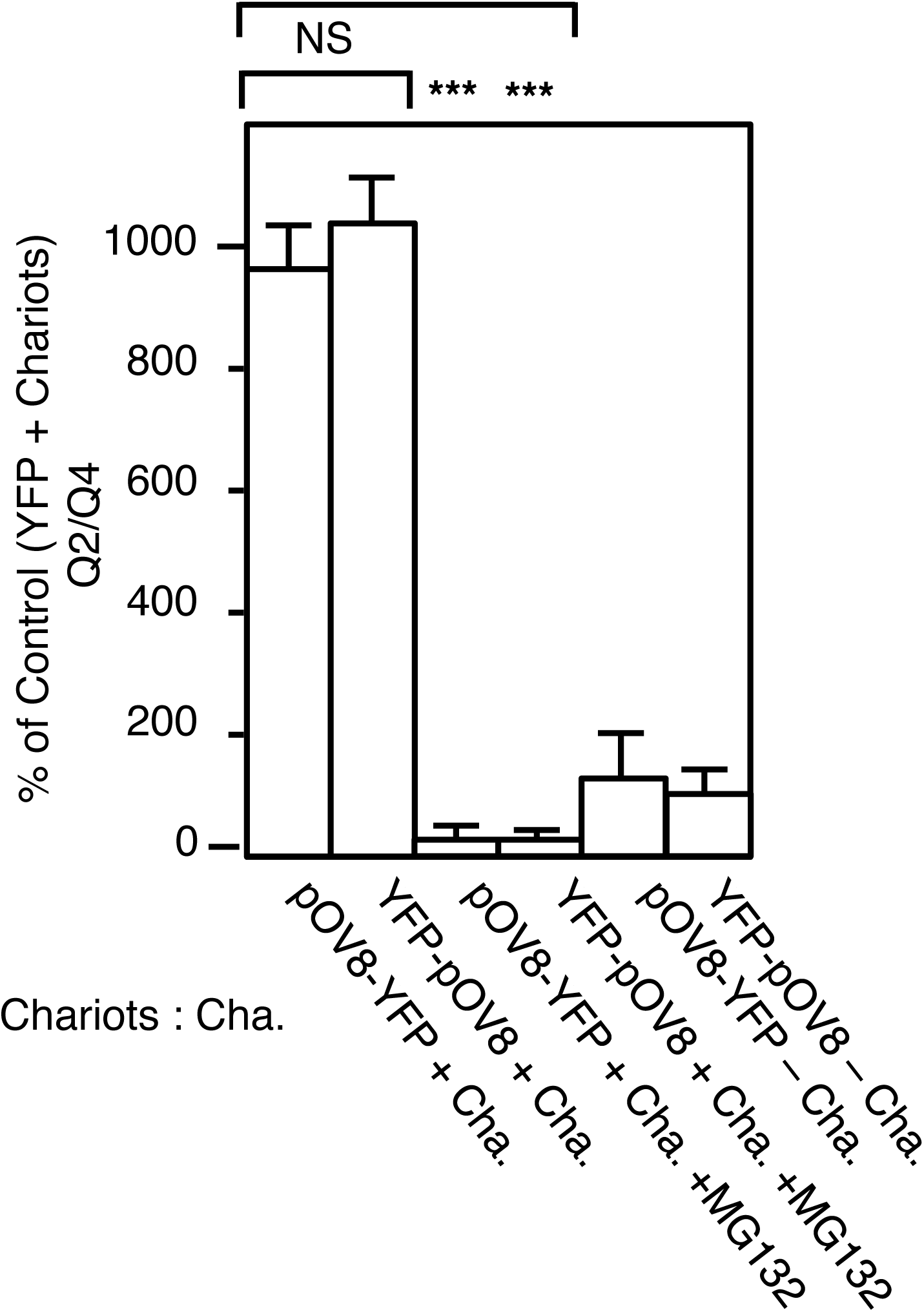
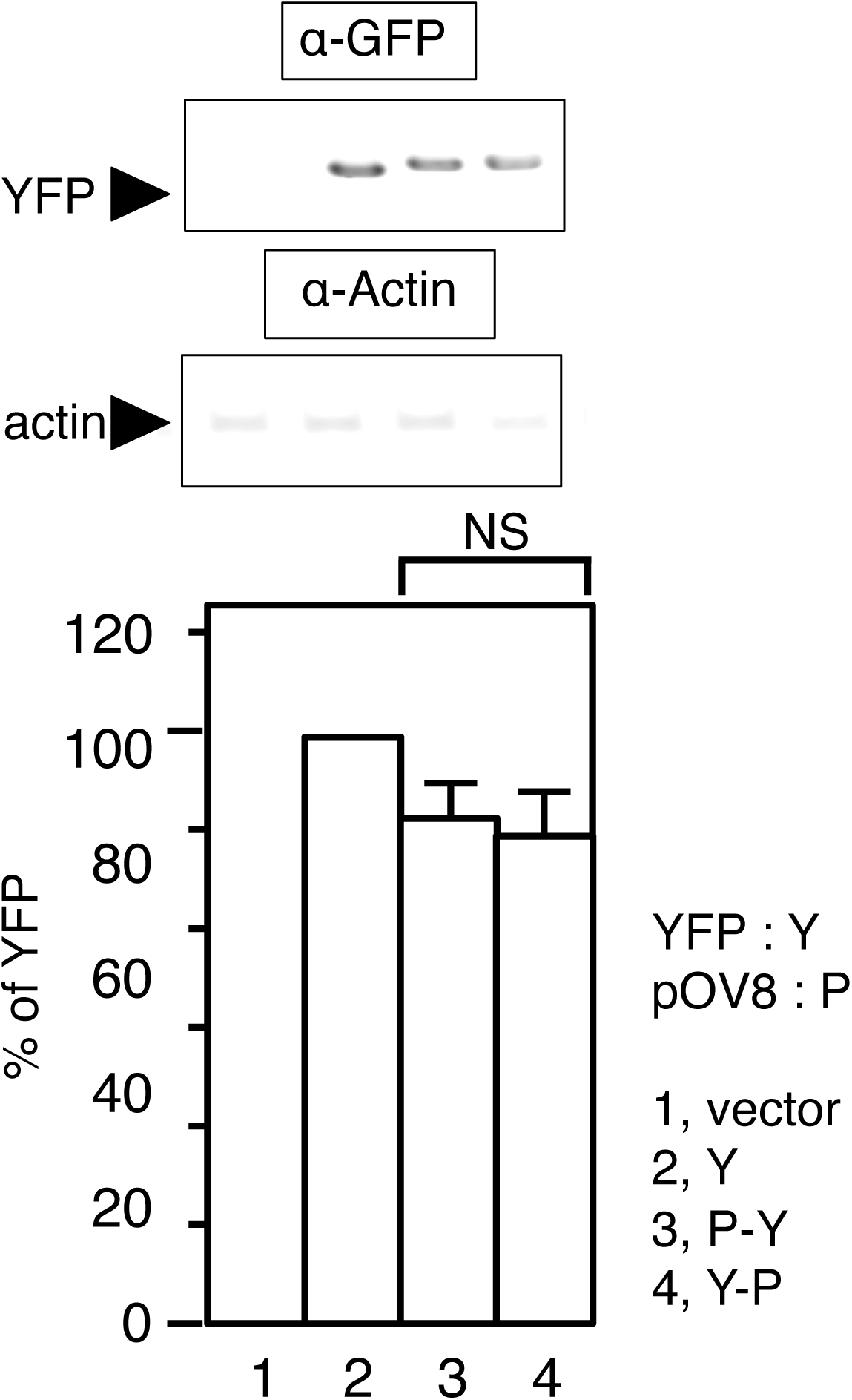
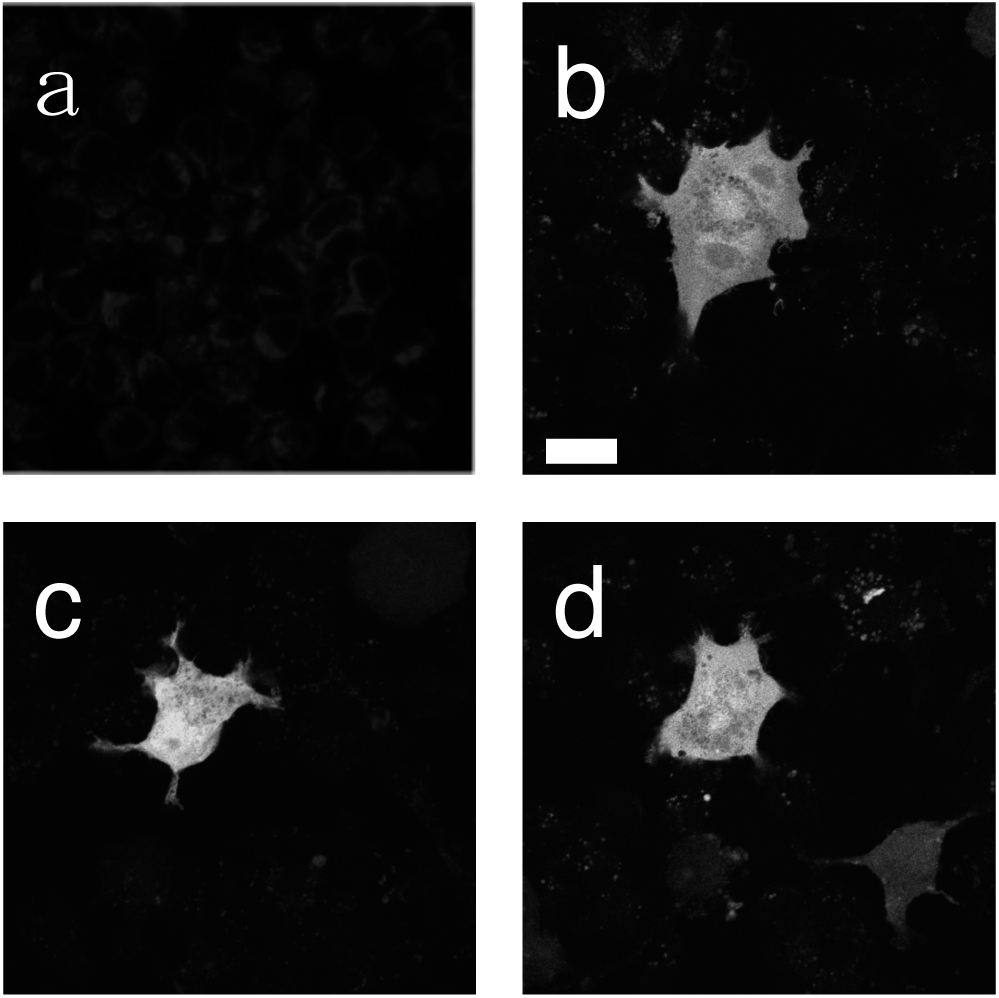
(A), Direct delivery of mature proteins into cells. The third of right-most; purified fluorescent proteins (YFP, pOV8YFP, and YFP-pOV8). DC2.4 cells were incubated with 5.6 μM of OVA (top of right-second and right-most), YFP (next of left-second), pOV8-YFP (next of right-second, third of left-second, and bottom of left-second), or YFP-pOV8 (next of right-most, third of right-second, and bottom of right-second) with (+ Chariot) or without Chariot^TM^ (only) and MG132. The fluorescent histograms of pOV8-YFP (pOV8-YFP + Chariot) and YFP-pOV8 (YFP-pOV8 + Chariot) were analyzed by color combine (pOV8-YFP/YFP-pOV8, third of left-most) as Fig. 1A. (B), Fluorescent intensities of mature proteins. Fluorescent intensities of the mature proteins (YFP, pOV8-YFP, and YFP-pOV8) were quantified by fluorometer and shown by relative intensities against fluorescent control protein (YFP). (C), Surface pOV8/Kb complex levels were quantified by Q2/Q4 and shown by relative amounts of fluorescent control protein (YFP). Student’s test was used to compare the pOV8-YFP by Chariot^TM^ sample against YFP-pOV8 by Chariot^TM^ sample, pOV8-YFP by Chariot^TM^ sample against pOV8-YFP by Chariot^TM^ sample with MG132, and YFP-pOV8 by Chariot^TM^ sample against YFP-pOV8 by Chariot^TM^ sample with MG132. (D), Localization of YFP (b), pOVA-YFP (c), and YFP-pOVA (d) in DC2.4. Intracellular localization of fluorescent proteins was detected by fluorescence of YFP. (a) as the control without fluorescent proteins. Bar, 10 μm. Data were shown as mean _±_ S.D. (error bars)., ***, p<0.01, NS, not significant, n=3.

### The predominant presentation of N-terminal antigenic peptides other than pOV8

Next, we examined whether the N-terminal predominance was observed with Ag peptides other than pOV8. Since we have no antibody other than 25-D1.16 to detect the peptide-MHC-I complex, we examined intra-molecular competition of pOV8 by Ag-peptides from GST or VSV. The VSV epitope has been shown to compete for K^b^ with pOV8 (Strehl et al., 2005). It is predicted that two independent Ag-peptides derived from GST would compete with pOV8 by database analysis (TABLE 1) (Johansen et al., 1997). As an Ag-peptide in the N-terminal of one protein was more efficiently complexed with MHC-I than the same Ag-peptide in the C-terminal of the same protein, when pOV8 binding was competed more effectively by N-terminally located GST (VSV) than C-terminally located GST (VSV), 25D-1.16 binding would be inhibited more efficiently by N-terminal GST (VSV) than C-terminal GST (VSV). We examined the effect of *in vitro* competition of pOV8 with GST derived peptides, GST48, GST111, and a peptide derived from VSV protein at first. DC2.4 cells were incubated with the constant concentration of peptides (50 μM), in which the decreasing concentrations of pOV8 (50-0 μM) with the increasing concentrations of competing Ag-peptides (VSV, GST48 or, GST111; 0-50 μM) were combined. In consequence, we compared pOV8 presentation under the following conditions; 50 μM of pOV8 and 0 μM of competitive peptide, 40 μM of pOV8 and 10 μM of competitive peptide, 25 μM of pOV8 and 25 μM of competitive peptide, 10 μM of pOV8 and 40 μM of competitive peptide, and 0 μM of pOV8 and 50 μM of competitive peptide (Fig. 3 *A* and *B*). Increasing amounts of each competitive peptide inhibited 25-D1.16 binding comparatively (Fig. 2 *A* and *B*), indicating that both GST48 and GST111 competed with pOV8 for K^b^ as VSV epitope. Then, we examined intra-molecular competition of pOV8 by GST or VSV, by comparing the pOV8 presentation between pOV8-YFP-GST (Fig. 2*C* upper-left, red) and pOV8-GST-YFP (Fig. 2*C* upper-left, green), between YFP-GST-pOV8 (Fig. 2*C* upper-right, red) and GST-YFP-pOV8 (Fig. 2*C* upper-right, green), between YFP-VSV-pOV8 (Fig. 2*D*, upper-left, red) and VSV-YFP-pOV8 (Fig. 2*D*, upper-middle, green). We again confirmed that expression levels (Fig. 2*C* bottom and 2*D* bottom) and localization of chimeric fluorescent proteins (Fig. 2*E*) were equivalent among each pair. Then we evaluated amounts of antigen presentation in addition to cell numbers; we quantified the amounts of pOV8/Kb presentations by the average fluorescence level of each histogram of Fig. 2*C* and *D* as described in Fig. 2*F*. All color combine histograms (Fig. 2*C* top, *D* upper-right) and quantification analysis (Fig. 2*G*) showed that N-terminally located GST (VSV) competed with pOV8 more efficiently than C-terminally located GST (VSV), indicating that the N-terminal GST (VSV) derived Ag-peptides were complexed with K^b^ more efficiently than the C-terminal GST (VSV) derived peptides (Fig. 2*C, D*, and *G*). These results suggest that the Ag-peptide located in the N-terminal of one protein is more effectively complexed with MHC-I compared with the same Ag-peptide in the C-terminal of the same protein.

**TABLE 1.** Calculated epitope scores on H2Kb (Over 15)

| Name |  |  | Name |  |  |
| --- | --- | --- | --- | --- | --- |
| Position | Sequence | Score | Position | Sequence | Score |
| OVA |  |  | GST |  |  |
| 258 | SIINFEKL | 25 | 48 | LGLEFPNL | 24 |
| 56 | KVVRFDKL | 22 | 111 | YSKDFETL | 24 |
| 177 | AEERYPIL | 22 | 169 | CLDAFPKL | 21 |
| 12 | NAIVFKGL | 22 | 152 | DFMLYDAL | 20 |
| 26 | CFDVFKEL | 20 | 3 | PILGYWKI | 18 |
| 96 | ENIFYCPI | 18 | 196 | SKYIAWPL | 17 |
| 28 | DVYSFSLA | 17 | 70 | AIIRYIAD | 15 |
| GFP |  |  | Azamigreen |  |  |
| 36 | GDATYGKL | 21 | 214 | AVARYSML | 23 |
| 38 | ATYGKLTL | 18 | 51 | LPFAYDIL | 21 |
| 5 | GEELFTGV | 17 | 201 | HDKDYNKV | 18 |
| 142 | LEYNYNSH | 17 | 203 | KDYNKVKL | 17 |
| VSV |  |  | 112 | GDCFFYDI | 16 |
| 52 | RGYVYQGL | 28 | 33 | NPYEGTQI | 15 |

Peptide used were: pOV8, SIINFEKL; VSV, RGYVYQGL; GST48, LGLEFPNL; GST111, YSKDFETL.
Scores were calculated by SYFPEITHI;
http://www.syfpeithi.de/scripts/MHCServer.dll/home.htm

### Kinetics of surface pOV8/Kb complex generation

To gain a more quantitative understanding of Ag-peptides generation, we examined the kinetics of pOV8/Kb complex formation in DC2.4 cells, which were transfected with either one of the pair of fluorescent chimeric proteins; pOV8-YFP and YFP-pOV8. After 12h, the cells were washed with acid (A.W.; <u>a</u>cid <u>w</u>ash) to remove preexisting K^b^ bound pOV8 on the cell surface, and the cells were then allowed to recover in normal medium for the indicated periods (Fig. 3*A*). Then we evaluated amounts of antigen presentation of Fig. 3 *B* and *C* as described in Fig. 2*F* (Fig. 3*D*). Though A.W. for 30 s removed more than 90% of the preexisting pOV8, a little difference remained between fluorescent histograms of the N-terminal pOV8 and the C-terminal pOV8 (Fig. 3*D* and Fig. 4*B* A.W.). Since elongated A.W. resulted in a marked increase for the rate of dead cells and hampered former experiments, we carried out following experiments by A.W. for 30 s. Because the expression of chimeric fluorescent proteins became plateau after 12 to 24 h and the amounts of pOV8/Kb complex rather reduced after 24 h in our assay system using transient transfectants of DC2.4, the recovery of a newly assembled pOV8/Kb complex on the cell surface was then assessed by 25-D1.16 for 12 h after A.W. (Fig. 3*A*). Both cells transfected with N-terminally located pOV8 and C-terminally located pOV8 generated the pOV8/Kb complex steadily and rapidly (Fig. 3*B, C*, and *D*) in these periods even 1 h from A.W. The rates of generation of pOV8/Kb complexes in YFP-pOV8 cells were about 50 % in compared with that of pOV8-YFP cells throughout our experiment, and the N-terminal predominance was eminent 4 h after the A.W. (Fig. 3*D*).

### The predominance of N-terminal Ag-peptide is dependent on the translation

We next investigated the efficiency of pOV8/Kb complex generation under the existence of several pharmacological inhibitors; α-amanitin for mRNA synthesis, anisomycin for peptidyl transfer, emetine for ribosomal translocation, amino acid analog, canavanine and azetidine acid, for protein folding, MG132 and lactacystin for proteasomes, and chloroquine for end/lysosome proteases. We measured the generation of newly synthesized pOV8/Kb complexes in standard medium for 6 h with several pharmacological inhibitors after the A.W. (Fig. 4*A*) and compared with the pOV8/Kb complex among N-terminal pOV8 and the C-terminal pOV8 (Fig. 4 *B, C*, and *D*). The generation of a pOV8/Kb complex was actively suppressed by inhibitors of proteasomes (Fig. 4*B, E*) but not with inhibitors of the end/lysosome proteases (Fig. 4*B, E*). Drugs inhibiting *de novo* protein synthesis also reduced the recoveries, but mainly influenced fluorescence histograms, measures of carrier proteins (Fig. 4*C*). Under these conditions, however, it was noteworthy that the predominance of N-terminus was largely diminished (Fig. 4*C, E*) without affecting pre-existing amounts of pOV8-YFP or YFP-pOV8 (Fig. 4*F*). On the contrary, drugs affecting protein folding thus promoting abnormal protein structures after translation, such as canavanine and azetidine acid (Fig. 4*D, E*), showed a significant increase of difference between the pair of histograms. These results indicate that the predominance of N-terminus is largely dependent upon *de novo* protein synthesis and degradations by proteasomes, but less dependent upon stabilities of proteins and degradations by end/lysosome proteases.

We then delivered purified mature fluorescent proteins (Fig. 5A third of the left-most) into cells directly. First, we introduced purified OVA into cells with or without MG132. 25-D1.16 staining (fraction Q1) was increased specifically in cells that introduced OVA (Fig. 5A top of the right-second) but decreased in cells with MG132 (Fig. 5A top of the right-most). The delivery of pOV8-YFP (Fig. 5A third of the left-second, red) and the delivery of YFP-pOV8 (Fig. 5A third of the right-second, green) showed no difference upon the generation of pOV8/Kb complex (Fig. 5A third of the right-most and Fig. 5C), indicating that when a pair of mature fluorescent proteins were delivered into cells directly, no predominance of N-terminus came about. These presentations were proteasomes-dependent because the simultaneous additions of MG132 with deliveries of fluorescent chimeric proteins strongly inhibited the generation of the pOV8/Kb complex (Fig. 5A bottom of left-second and bottom of right-second), but they also showed a marked reduction for the rate of living cells. Besides, these presentations were not the results from cross-presentation because the small amounts of the pOV8/Kb complex were detected without the delivery reagent (Fig. 5A next to the right-second, next to the right-most, and C). Again, we checked that there were no differences in amounts of delivered proteins (Fig. 5D) nor localization of these proteins (Fig. 5E).

### The effects of IFN-γ upon the predominant presentation of N-terminal antigenic peptides

We last investigated the effect of IFN-γ, which enhances expression of MHC-I and substantially alters the cleavage sites of substrates and therefore dramatically improves Ag presentation (Rammensee et al., 1999), upon the predominance of the N-terminal pOV8 (Fig. 6*A*). We quantified the amounts of pOV8/Kb presentations by the average fluorescence level of each histogram of Fig. 6*A* as Fig. 2*F*. The addition of IFN-γ enhanced the amount of pOV8/Kb about 2.2- and 3.1-fold in our experimental conditions from pOV8-YFP and YFP-pOV8 respectively by the whole histogram (Fig. 6*B* left, TP_20-10000_ of IFN-γ+/ IFN-γ- for pOV8-YFP and YFP-pOV8) and 1.9- and 2.8-fold by 10^3^ fluorescent units of YFP (Fig. 6*B* left, TP_20-1000_ of IFN-γ+/ IFN-γ- for pOV8-YFP and YFP-pOV8). N-terminal pOV8 complexed with K^b^ was about 1.9-folds high compared with C-terminal pOV8 by the whole histogram (Fig. 6*B* right, TP_20-10000_ of pOV8-YFP/YFP-pOV8 for IFN-γ-) or 3.1-folds by 10^3^ fluorescent units (Fig. 6*B* right, TP_20-1000_ of pOV8-YFP/YFP-pOV8 for IFN-γ-) in absence of IFN-γ, and 2.1-folds high by the whole histogram (Fig. 6*B* right, TP_20-10000_ of pOV8-YFP/YFP-pOV8 for IFN-γ+) or 2.8-folds by 10^3^ fluorescent units (Fig. 6*B* right, TP_20-1000_ of pOV8-YFP/YFP-pOV8 for IFN-γ+) in presence of IFN-γ. These results indicate that pOV8 located in the N-terminal of one protein was more efficiently complexed with MHC-I compared with pOV8 located in the C-terminal of the same protein. And that, this predominance was independent of the presence of IFN-γ (Fig. 6*A, B*).

**Figure 6.**
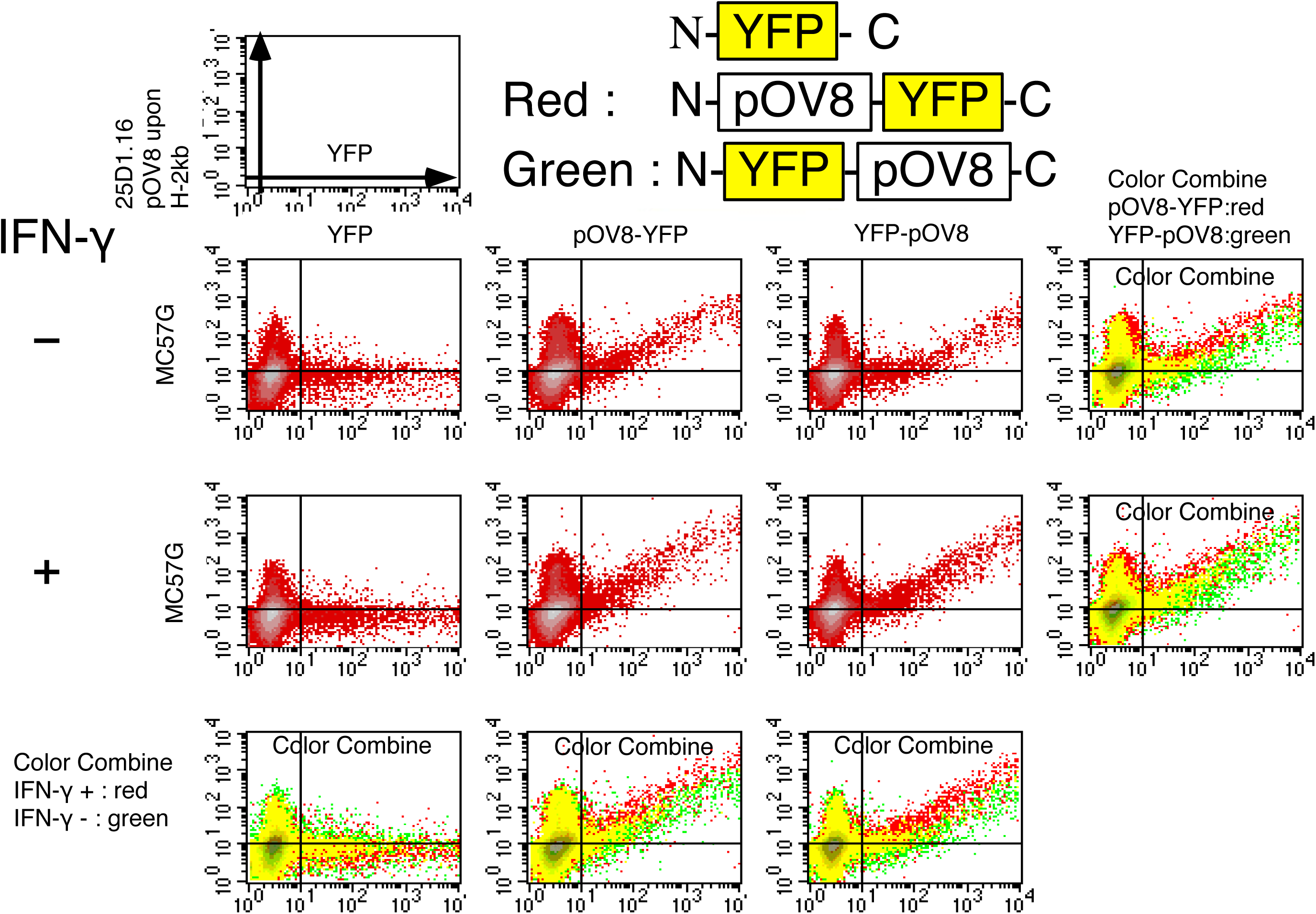
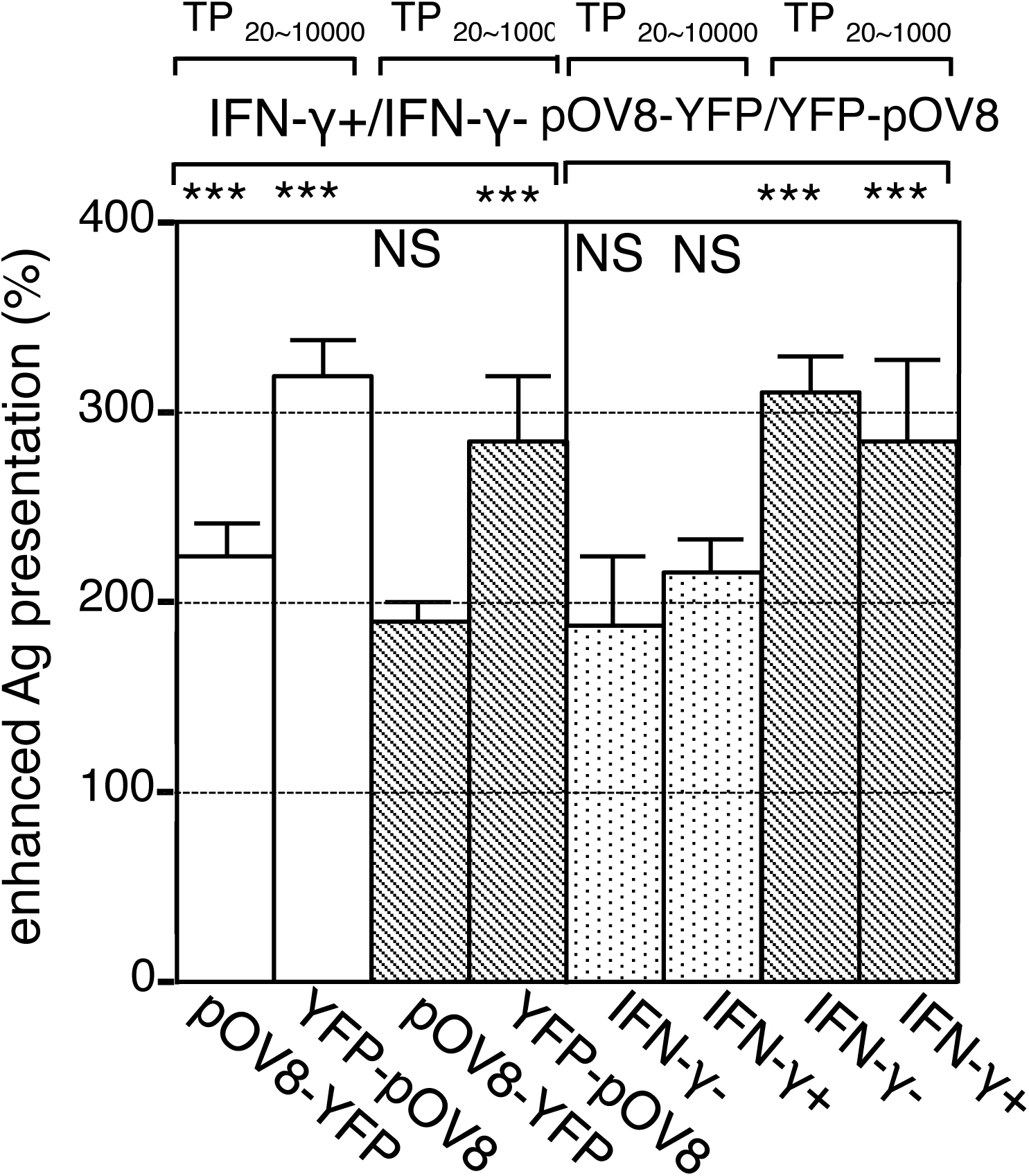
The effects of IFN-γupon pOV8/Kb complex. (A), pOV8/Kb complex generation from MC57G cells transfected with pOV8-YFP or YFP-pOV8 with or without 100 units/ml of IFN-γ. The color combine analyses of the right side were designated as above. The lowest lane was a combination of above histograms coloring with IFN-γ as red and without as green. (B), Quantification of enhanced presentation of pOV8/Kb complex by treatments of IFN-γ and the N-terminus location shown in Fig2 F. Student’s test was used to compare (IFN-γ+/IFN-γ-, MC57G, pOV8-YFP) sample with (IFN-γ+/IFN-γ-, MC57G, YFP-pOV8) sample, (pOV8-YFP/YFP-pOV8, MC57G, IFN-γ+) sample, and (pOV8-YFP/YFP-pOV8, MC57G, IFN-γ-) sample in both Whole samples and ∼10^3^ fluorescent unit respectively. Data were shown as mean _±_ S.D. (error bars). ***, p<0.01, **,p<0.1, NS, not significant, n=3.

## Discussion

In this report, we showed a novel rule for the direct presentation; an Ag peptide located in the N-terminus of a protein is more efficiently complexed with MHC-I compared with the same Ag-peptide located in the C-terminus of the protein (Fig. 1). This predominance was independent of cell types (Fig. 1*A*), proteins harboring the Ag-peptide (Fig. 1*A, G*), the kind of Ag-peptides (Fig. 2), and the immunomodulation by IFN-γ (Fig. 6A). Since in series of our experiments the antigen presentation efficiencies were compared N-terminally tagged fluorescent proteins and C-terminally tagged proteins based on the fluorescence intensities, we showed every biological property of the pair of fluorescent protein shown below was equivalent: the fluorescent intensity (Fig. 5*B*), the transcriptions (Fig. 1*C*), the amounts (Fig. 1*D*), the cellular localization (Fig. 1*H*), and correlation of the fluorescent intensity and the expression level (Fig. 1*D*). In addition, 4-6 h was enough to bring about the N-terminal predominance (Fig. 3*B, C*, and *D*), amounts of pOV8-YFP and YFP-pOV8 were equivalents after 6 h of treatments by Cycloheximide (Fig. 1*E*) and α-amanitin (Fig. 4*F*), indicating that the stability of the pair of fluorescent protein was equivalent in this meanwhile. Throughout these experiments, we can find a significant population of pOV8/Kb-positive cells that were not fluorescent. These cells, which were pOV8/Kb-positive but did not show any fluorescence, were the results of non-specific binding of 25D1.16 antibody upon cell surfaces, because these populations were detectable in control cells without pOV8 (Fig. 1*A*, YFP and Fig. 5*A* Chariot only), did not decrease after acids wash (Fig. 3*B* and *C*).

Although we were able to examine the rule only for K^b^, because of the limited availability of antibody for peptide-MHC-I complex, our results strongly suggest that this rule is conserved in the process of the direct presentation. It should be noted that the effects of the N-terminal predominance were comparable with the enhanced MHC-I presentation by IFN-γ (Fig. 6*A, B*). We first quantified the amounts of the pOV8/Kb complex from the whole histogram up to 10^4^ fluorescent units of the horizontal axis-YFP (Fig. 6*B*, left). Thousands of fluorescent units for vertical axis-25D1.16 correspond to full surface MHC-I as judged from the addition of excess amounts of the pOV8 peptide, indicating that the most of the surface MHC-I (10^4^-10^5^ MHC-I molecules) were occupied by pOV8. This saturation of MHC-I differ from physiological conditions, and thus pOV8-YFP showed a small difference with or without IFN-γ compared with YFP-pOV8 (Fig. 6*B*, left). It is noteworthy that 10^1^ fluorescent units of vertical axis-25D1.16 correspond to no pOV8/Kb complex (Fig.1 *A*, YFP histogram). We re-quantified and compared the amounts of pOV8/Kb from a restricted area of each histogram up to 10^3^ fluorescent units of the horizontal axis-YFP (Fig. 6*B* right). The resultant re-quantification showed that the effect of N-terminal predominance was comparable with the effect of IFN-γ (Fig. 6*B* right). Since CTL can be triggered by a small number of complexes (Wherry et al., 1999), these results suggest that the effect of N-terminal predominance plays a pivotal role under physiological conditions.

What is the molecular mechanism determining the N-terminal predominance? Ag-peptides are for the most part provided from two sources; retirees and DRiPs (Lelouard et al., 2002; Lelouard et al., 2004; Pierre, 2005; Yewdell et al., 1996). Besides, a subset of the nascent polypeptide, which is directly delivered to the 20S proteasome, also provides Ag-peptides (Starck et al., 2016; Voo et al., 2004; Yewdell and Nicchitta, 2006). Retirees are once matured proteins, which completed their lifetimes or damaged by environmental stress. In contrast, DRiPs are nascent proteins, which is terminated before completing their synthesis. A large proportion of newly translated proteins (up to 30 %) are degraded as DRiPs before maturation. DRiPs are further divided into premature termination products (tDRiPs) and misfolded full-length products (mDRiPs) (Yewdell et al., 1996). Since all proteins start their translation from the N-terminal methionine and complete at the C-terminal termination codons, while retirees and mDRiPs include equivalent amounts of N-terminus and C-terminus Ag-peptides, most of the tDRiPs contain N-terminal Ag-peptides without C-terminal Ag-peptides. Because most tDRiPs only provides N-terminal Ag-peptides, the predominance of N-terminal peptides is likely derived from tDRiPs. In our experimental conditions, the N-terminal predominance was dependent upon degradation by proteasomes (Fig. 4*B, E*) and *de novo* protein synthesis (Fig. 4*C, E*), strongly indicating that the N-terminal predominance was dependent upon DRiPs. It is also noteworthy that the N-terminal predominance was increased to some extent by amino acid analogs, canavanine and azetidine acid, which inhibit protein folding and results in an increase of mDRiPs, suggesting that the N-terminal predominance was dependent more upon tDRiPs and less upon mDRiPs (Fig. 4*D* right two columns and *E*). Also, the forced introduction of folded proteins showed little difference between N-terminal epitope and C-terminal epitope, indicating that the N-terminal predominance was dependent upon the *de novo* protein synthesis (Fig. 5*A, C*). All the above results were showing that the N-terminal predominance is results of tDRiPs and that this rule would be a general rule for the direct presentation.

As noted previously, GFP variants produce mortally shows fragmentation to create DRiPs (Wei et al., 2015). These Triton X-100-insoluble proteasome substrates were hardly detectable in our experimental conditions. It might be because these by-products were results of chromophore rearrangements, transfected cells are kept in the dark condition in the course of experiments (Barondeau et al., 2006; Remington, 2006). Also, though these fragmentations would also contribute Ag-presentation in our assay system, they would not bring about the N-terminal predominance, because these fragmentations were classified mDRiPs. As noted above the contribution of mDRiPs upon N-terminal predominance was small enough compared with tDRiPs. Furthermore, we also observed the N-terminal predominance with structurally distinct fluorescent proteins, Azamigreen (Fig. 1*G*). These results strongly deny the N-terminal predominance was the results of specific fragmentations by GFP variants.

DC can present exogenous Ag upon MHC-I (Imai et al., 2205; Villadangos et al., 2007), which is called cross-presentation (CP). The CP is independent of protein synthesis in DCs (Donohue et al., 2006; Palliser et al., 2005) and thus unrelated to N-terminal predominance. In cancer cells or virally infected cells, protein translation rates are high enough to produce excess amounts of unfolded proteins for the protein degradations machinery then these cells produce unfolded proteins or aggregated proteins. Since unfolded proteins interact with several kinds of molecular chaperones and molecular chaperones are known to enhance the efficiency of CP (Binder and Srivastava, 2005; Flechtner et al., 2006; Kunisawa and Shastri, 2006). Aggregated proteins were incorporated into autophagosomal membranes and released from the cell surface by unconventional protein secretion. These secretory fractions were called DRibbles (DRibbles; <u>D</u>efective <u>Rib</u>osomal products-containing autophagosome-rich <u>ble</u>bs). DRibbles were the more effective source for CP in DC compared with cell lysate and soluble antigens (Hampton, 2002; Wiertzn et al., 1996). Since those unfolded proteins and aggregated proteins inevitably contain tDRiPs, it might be possible that the N-terminal predominance might be detectable in CP, because CTL attack target cells by recognizing peptides with MHC-I, the peptide selection between DCs and target cells would be key for the effective cell-mediated immunity. Because only small numbers of peptide-MHC-I complex per DC is sufficient for activating CD8+ T cells (Palliser et al., 2005), the N-terminus predominance might play important roles in cell-mediated immunity.

Recent reports have demonstrated that no significant preference was observed for N-terminally located antigenic peptides. Either by the *in silico* analysis exploiting the available data in the Immune Epitope Database and Analysis Resource (Hassan et al., 2013; Kim et al., 2013) or by *in vitro* analysis using dynamic stable isotope labeling by amino acids in cell culture (Bourdetsky et al., 2014; Hassan et al., 2013) showed no positional skew. It is possible to describe our observations as artifacts by several assumptions; it might be plausible that our observations were results of vector-dependent mistranslations, such as the premature translation termination (tDRiPs), impaired stability (mDRiPs), or the downstream initiation (Berglund et al., 2007). Since recent researches strongly designate that DRiPs are one of the main sources of antigenic peptides (Antón et al., 2014), though the N-terminal predominance was overestimated in our assay system, it is also possible that this rule has the definite possibility as the real cellular phenomenon. On the contrary, the downstream initiation rather induces the C-terminal predominance, even if that was the case, it rather reduced the N-terminal predominance.

Nevertheless, it is also likely that large-scale studies harbor their experimental system-specific errors. *In Sirico* analysis, it is likely that CTL epitopes for viral Ags are largely dependent upon the affinity against MHC I and less dependent upon the location of the Ag-peptide. In SILAC experiments, antigenic peptides were purified by a specific antibody against MHC I (Bourdetsky et al., 2014). Through purifications, antigenic peptides with lower affinity against MHC I might be lost. In addition, there were rather restricted numbers of antigenic peptides, which were both available for the ratio between the rates of synthesis of themselves and of their source proteins, because the proteome analysis has lower sensitivity against proteins with low expression level or proteins with lower stability. In consequence, analyzed antigenic peptide tend to be derived from abundant proteins or to show higher affinity with MHC I. Since the quantity of the precursor peptide and the affinity of the precursor peptide against MHC I are the most fundamental qualification of antigenic peptides upon MHC I, these antigenic peptides show a smaller effects with the positions of their source. Hence the positional skew would be under quantified in SILAC experiments. Large-scale experiments are effective methods to overlook the cellular phenomenon, but these experiments can’t directly compare the positional skew of the same antigenic peptide derived from the equivalent source proteins. In contrast, a cell-based assay system utilizing specific probes shows higher distinctiveness against restricted cellular phenomena. Both methods are indispensable to elucidate complicated cellular phenomenon.

In this study, we found a novel rule for the direct presentation, the N-terminal predominance of Ag-peptides. In cancer immunotherapy, it is very difficult to find out effective Ag-peptides, which induce T cell responses against cancer. Since cancer is derived from self, CD8^+^ T cells strongly reactivate with self-peptides were excluded in the thymus or fell in anergy (Janeway et al., 2001). Thus it is likely that CTLs reactivated with natural cancer derived peptide-MHC-I complexes are likely of low affinity against the complexes supplemented by the number of the complexes as higher avidity. The higher efficiency of the N-terminal Ag-peptide presentation results in higher avidity and might show a new criterion for finding out effective cancer Ag-peptides. In addition, the introduction of cancer-specific Ag-peptide as a fusion protein with a carrier protein into cancer to induce T cell responses is one of the strategies for cancer immunotherapy (Lei et al., 2015; Petrovsky et al., 2004; Rappuoli et al., 2011). The N-terminal predominance might provide pivotal information upon the configurations of recombinant proteins. An important question is whether N-terminal predominance has any physiological roles in the immune system. However, some clinical studies still indicated that N-terminally located Ag-peptides elicit stronger immune responses compared with C-terminally located Ag-peptides (Kawashima et al., 1998; Ma et al., 2004; Monji et al., 2004). These results suggest that the N-terminal predominance might play certain roles in our immune system.

## Materials and Methods

### Cells and cultures

DC2.4, a DC line (31), was provided by Dr. K. L. Rock (Dana-Farber Cancer Institute). EL4, a lymphoma line, was our laboratory stock. MC57G, a fibrosarcoma cell line, was purchased from ATCC. Cells were cultured in RPMI-1640 (SIGMA) supplemented with 2 mM L-glutamine, 1 mM sodium pyruvate, 0.1 mM non-essential amino acid, 100 units ml^-1^ penicillin-streptomycin, 55 μM 2-mercaptoethanol, 10 mM Hepes (pH 7.5) and 10% FCS or DMEM (Gibco) supplemented with 10% FCS with or without 100 units of IFN-γ (Becton Dickinson) at 37°C in 5% CO_2_ unless otherwise indicated. Polymyxin B (50 μg/ml) was present throughout in cell cultures. *Plasmids* The pOV8 epitope (SIINFEKL) with each five flanking amino acid in its N- and C-terminus (LEQLESIINFEKLTEWTS) was created using convergent primers, 5′-GCGTGATCACTTGAGCAGCTTGAGAGTATAATCAACTTTGAAAAACTGAC-3′ and 5′-CGCAGATCTGGATCCACTGGTCCATTCAGTCAGTTTTTCAAAGTTGAT-3′. The created fragment harbored N-terminal *Bcl* I site, C-terminal *Bam* HI site and C-terminal *Bgl* II site, and was sub-cloned into the pT7 vector to produce pT7-pOV8. Then, the *Sma* I site of pT7-pOV8 was changed to *Bgl* II site with ten mer *Bgl* II linker (GCAGATCTCG, TAKARA) to produce pT7-Bgl II-pOV8. VSV epitope (RGYVYQGL) with each five flanking amino acid in its N- and C- terminus (LEQLERGYVYQGLTEWTS) was created using convergent primers, 5′-GCGAGATCTATGCTTGAGCAGCTTGAGCGTGGTTATGTTTATCAAGGTTT-3′ and 5′-CGCCTCGAGGGATCCACTGGTCCATTCAGTTAAACCTTGATAAACATAACC-3′. The created fragment was used as pOV8. The complete YFP, OVA, GST, Azamigreen coding region was amplified by PCR from appropriate plasmids using convergent primers; YFP with 5′-GCGAGATCTATGGTGAGCAAGGGCGAG-3′ and 5′-CGCGGATCCGAGTCCGGACTTGTACAGCTC-3′, OVA with 5′-GCGAGATCTATGGGCTCCATCGGT-3′ and 5′-CGCGGATCCTAGCTAGCAGGGGAAACACATCTG-3′ Azamigreen with 5′-GCGAGATCTATGGTGAGTGTGATTAAA-3′ and GST with 5′-GCGAGATCTATGTCCCCTATACTAGGT-3′ and 5′-CGCAGATCTTCAGGATCCACGCGGAACCAGATCC-3′. All created fragments harbor N-terminal *Bgl* II site and C-terminal *Bam* HI site and were subcloned into the pT7 vector to produce, pT7-YFP, pT7-OVA, pT7-Azamigreen, and pT7-GST. Because *Bam* HI site, *Bgl* II site, and *Bcl* I site are all compatible with each other, we inserted YFP fragments from *Bgl* II site to *Bam* HI site of pT7-YFP into *Bgl* II site of pT7-GST and pT7-OVA to produce pT7-YFP-GST and pT7-YFP-OVA or into *Bam* HI site of pT7-GST and pT7-OVA to produce pT7-GST-YFP and pT7-OVA-YFP. Above fragments were then inserted into the pEF/myc/cyto (Invitrogen) vector with some modifications. The *Pst* I site of pEF/myc/cyto was changed to *Bgl* II site with ten mer *Bgl* II linker (above) after blunting. Then the fragment between *Bgl* II site and *Bcl* I site were converted using convergent primers to produce pEF-Bgl II; 5′-GATCTTAACTAGCTGATCAGATATCG-3′ and 5′-GATCTTAACTAGCTGATCAGATATCG-3′. To produce expression vectors used in this study, we inserted fragments described above into pEF-Bgl II vector in order of the indicated name. To produce pEF-YFP, pEF-OVA, pEF-YFP-OVA, pEF-OVA-YFP, pEF-Azamigreen, pEF-GST-YFP, and pEF-YFP-GST, YFP, OVA, YFP-OVA, OVA-YFP, Azamigreen, GST-YFP, and YFP-GST fragments of *Bgl* II site to *Bam* HI site were inserted into *Bgl* II site of pEF-Bgl II, respectively. To produce pEF-pOV8, the pOV8 fragment of *Bcl* I site to *Bgl* II site was inserted into pEF-Bgl II. Then YFP, Azamigreen, GST-YFP, and YFP-GST fragments of *Bgl* II site to *Bam* HI site were inserted into *Bam* HI and *Bgl* II site of pEF-pOV8 to produce pEF-pOV8-YFP, pEF-pOV8-Azamigreen, pEF-pOV8-GST-YFP, pEF-pOV8-YFP-GST, respectively. To produce pEF-Bgl II-pOV8, pOV8 fragment from *Bgl* II site to *Bam* HI from pT7-Bgl II-pOV8 was inserted into pEF-Bgl II. Then YFP, Azamigreen, GST-YFP, and YFP-GST fragments of *Bgl* II site to *Bam* HI site were inserted into *Bgl* II and *Bcl* I site of pEF-Bgl II-pOV8 to produce pEF-YFP-pOV8, pEF-Azamigreen-pOV8, pEF-GST-YFP-pOV8, pEF-YFP-GST-pOV8, respectively. For expression in *E.coli,* pET30a-YFP, pET30a-pOV8-YFP, and pET30a-YFP-pOV8 were produced as above series of pEF vectors. To produce pT7-pOV8-YFP, the YFP fragments from *Bgl* II site to *Bam* HI site from pT7-YFP were inserted *Bam* HI site of pT7-pOV8 to produce pET30a-YFP. To produce pT7-YFP-pOV8, the pOV8 fragments from *Bcl* I site to *Bam* HI site from pT7-pOV8 were inserted *Bam* HI site of pT7-YFP to produce pT7-YFP-pOV8. The YFP fragments of *Bgl* II site to *Bam* HI, the pOV8-YFP fragments of *Bcl* I site to *Bam* HI site, and the YFP-pOV8 fragments of *Bgl* II site to *Bam* HI site, were inserted *Bam* HI site of pET30a (Qiagen) to produce pET30a-YFP, pET30a-pOV8-YFP, and pET30a-YFP-pOV8.

### Transfection

All plasmids used in this study were purified by EndoFree Plasmid Purification system (Qiagen) according to the supplier’s protocol to remove endotoxin. Cells were transfected with these plasmids using Superfect Transfection Reagent (Qiagen).

### Antibodies and reagents

Anti-GFP (rabbit for Western blotting; MBL, mouse clone E4 and RQ2 for in cell staining; MBL), Anti-Azamigreen (rabbit; MBL), Anti-actin (mouse; Millipore), and 25-D1.16 were used in this study. All peptides used in peptide loading experiments were obtained from Peptide Institute Inc. CHX, MG132, Lactacystin, chloroquine, α-amanitin, Anisomycin, Emetine, L-Canavanine, Azetidine-3-Carboxylic Acid, purified ovalbumin (OVA) were purchased from Sigma Chemical Co. Protein concentration was determined using BCA protein assay reagent (Pierce Chemical Co.) with bovine serum albumin (Sigma) as the standard.

### Antigen presentation assay

pOV8/Kb levels were determined by incubating cells for one h on ice with the 25-D1.16 antibody followed by one h on ice with secondary goat anti-mouse antibody conjugated to Alexa Fluor 647 (Molecular Probes). Cellular fluorescent proteins and Alexa Fluor 647 levels were determined on a FACS Calibur cytofluorograph (Beckton Dickinson) using CellQuest (Beckton Dickinson) software (Fig. 1*A, F, G*, 2*A, C, D, F*, 3*B, C*, 4*B, C, D*, 5*A*, and 6*A*).

### In cell staining

Cells were washed twice with FACS buffer (2.68 mM KCl, 1.47 mM KH_2_PO_4_, 136.89 mM NaCl, 8.10 mM Na_2_HPO_4_, 1% BSA) and fixed by 2% formaldehyde in FACS buffer at 25°C for twenty min. Fixed cells were washed twice by FACS buffer and permeabilized by 2% saponin in FACS buffer at 25°C for ten min. The amounts of actin (YFP) were determined by incubating cells for one h on ice with the ant-actin antibody (anti-GFP antibody clone E4 or RQ2) followed by one h on ice with secondary goat anti-mouse antibody conjugated to Alexa Fluor 647 (Molecular Probes). Cellular fluorescent proteins and Alexa Fluor 647 levels were determined on a FACSCanto II (Beckton Dickinson) using BD FACSDiva^TM^ (Beckton Dickinson) software (Fig. 1*B*).

### RT-PCR

Total cellular RNA was isolated by the NucleoSpin^®^ RNA II kit (Macherey-Nagel), according to the manufacturer’s recommendations. RNA (500 ng) was reverse- transcribed to cDNA (BcaBEST™ RNA PCR Kit Ver. 1.1), subjected to real-time PCR. The expression level of each gene is normalized to ubiquitin. The oligonucleotides used are described below.

Ubiquitin Forward 5′-GCGAGATCTATGCAAATCTTTGTGAAAAC-3′

Ubiquitin Reverse 5′-CGCGGATCCCCCACCTCTGAGGCGAAGGACCA-3′

YFP Forward 5′-GCGAGATCTATGGTGAGCAAGGGCGAG-3′

YFP Reverse 5′-CGCGGATCCGAGTCCGGACTTGTACAGCTC-3′

### Western blotting

Cells were washed twice with PBS and solubilized in TNE (20 mM Tris-HCl pH 7.4, 150 mM NaCl, 0.5 M EDTA, 1% Nonidet P-40) with protease inhibitor cocktails (Sigma), cell debris were removed by centrifugation (16,000 X g, 5 min, 4°C). In case to detect fragmented GFP, cells were solubilized in SDS extraction buffer (1% SDS, 50 mM Tris-HCl (pH 7.4), 5 mM EDTA, 15 units/ml DNase) with protease inhibitor cocktails (Sigma), cell debris were removed by centrifugation (16,000 X g, 5 min, 4°C). Cell lysates were subjected to SDS-PAGE transferred onto PVDF membrane. Then the blot was developed with the indicated primary antibodies, horseradish peroxidase conjugated secondary antibodies and the chemiluminescent substrate.

### Microscopy

Images were obtained using A1 ver. 4.60 (Nikon, Tokyo, Japan) laser confocal microscopy.

### Peptide loading experiments

DC2.4 cells were incubated for one h at 37°C in medium with indicated peptides. After washed twice with PBS, the amounts of the pOV8/Kb complex were determined by

### Acid wash recovery assay

DC2.4 cells were washed with PBS and exposed to 0.131 M sodium citrate and 0.066 M sodium phosphate (pH 3.10). After 30 sec incubation, cells were washed twice with PBS and once with a medium. Then cells were grown in the medium for the indicated time with or without inhibitors.

### Quantification of the amounts of pOV8/Kb complexes

*s*The total amount pOV8/Kb (TP) of each histogram was quantified by adding up each average fluorescents unit for the vertical axis-25D1.16 (APk) multiplying cell numbers of the corresponding fluorescents unit for horizontal axis-YFP (CNk);

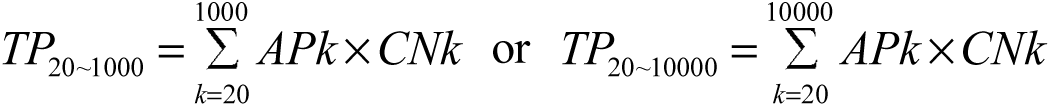

APks were worked out by the approximation curve of each histogram plotted horizontal axis-YFP against vertical axis-25D1.16 as Fig. 2*F*. Since cells with low fluorescents unit of YFP contained cells without antigenic proteins, AP_1∼10_ was left out. CNks were calculated by each histogram plotted YFP against cell number as Fig. 2*F* right-most. Without any comments, TP_20∼10000_ were indicated as quantifications (Fig. 2*G*, 3*D*, 4*E*).

### Purification of recombinant proteins

His-tag purified recombinant proteins were expressed in *E.coli* and purified by Ni^+^-NTA-agarose column (Qiagen). Then, contaminated endotoxin was removed by Detoxi-Gel (PIERCE Biotechnology, Inc.), twice according to the supplier’s protocol.

### Direct delivery of purified proteins

For direct delivery of purified proteins into cells, we used the amphipathic peptide carrier system, called Chariots^TM^ (AnaSpec, Inc.) according to the supplier’s protocol.

### Fluorescent Intensity

The fluorescent intensities of purified proteins were obtained using Perkin Elmer 1420 multi-label counter ARVO^TM^ MX fluorescent spectrum meter, CW-lamp filter name F485, CW-lamp filter slot A5, CW-lamp energy 40000, emission filter name F535 nm, emission filter slot A5, and measurement time for 1.0 S.

## Acknowledgments

This work was supported in part, a Grant-in-Aid for Scientific Research on Priority Areas (C) (17570164), a National Grant-in-Aid for the Establishment of a High-Tech Research Center in a private University, the Takasaki University of Health and Welfare, a Grant for the promotion of the advancement of education and research in graduate schools, and a Scientific Frontier Research Grant from the Ministry of Education, Culture, Sports, Science and Technology, Japan.

## Author contribution

J.I., M.O., E.K., Y.Y., M.M., S.K., and T.S. conducted the experiments; J.I designated the experiments; and J.I. and T.S. wrote this paper.

## Coflict of Interests

The authors declare no competing interests.

